# How many mammal species are there now? Updates and trends in taxonomic, nomenclatural, and geographic knowledge

**DOI:** 10.1101/2025.02.27.640393

**Authors:** Connor J. Burgin, Jelle S. Zijlstra, Madeleine A. Becker, Heru Handika, Jesse M. Alston, Jane Widness, Schuyler Liphardt, David G. Huckaby, Nathan S. Upham

## Abstract

The Mammal Diversity Database (MDD) is an open-access resource providing up-to-date taxonomic, nomenclatural, and geographic data for global mammal species. Since its launch in 2018, the MDD has transformed the traditionally static process of updating mammalian taxonomy into regular online releases reflecting the latest published research. To build on this foundation, we here present version 2.0 of the MDD (MDD2), which catalogues 6,759 living and recently extinct mammal species, representing net increases of 4.1% and 24.8% over MDD version 1.0 and *Mammal Species of the World*, 3rd edition (MSW3), respectively. Additionally, we identify a net increase of 68.8% (+2,754; 3,149 splits + de novo, 395 lumps) species since 1980 at a rate of ∼65 species/year based on past totals from 14 mammalian compendia, leading to projections of ∼7,084 species by 2030 and ∼8,382 by 2050 if these trends continue. Key updates in MDD2 include: (i) codings of US state, country, continent, and biogeographic realm geographic categories for each species; (ii) a comprehensive nomenclatural dataset for 50,230 valid and synonymous species-rank names, curated with type locality and specimen information for the first time; and (iii) integration between the MDD and the databases Hesperomys and Batnames for greater data accuracy and completeness. These updates bridge critical gaps in the taxonomic and nomenclatural information needed for ongoing revisions and assessments of mammalian species diversity. Using these data, we evaluate temporal and geographic trends over the past 267 years, identifying four major time periods of change in mammalian taxonomy and nomenclature: (i) the initial monographic description of traditionally charismatic species (1758–1880); (ii) the peak of descriptive taxonomy, describing subspecies, and publishing in journals (1881–1939); (iii) the shift toward revisionary taxonomy and polytypic species (1940– 1999); and (iv) the current technology-driven period of integrative revisionary taxonomy (2000– present). Geographically, new species recognition since MSW3 has been concentrated in equatorial, mountainous, and island regions, highlighting areas of high mammal endemism (e.g., Madagascar, Philippines, Andes, East Africa, Himalayas, Atlantic Forests). However, gaps in 21st century taxonomic activity are identified in West and Central Africa, India, and some parts of Indonesia. Currently lagging conservation assessments are alarming, with 25% of the MDD2-recognized mammal species allocated to the ‘understudied’ conservation threat categories of Data Deficient (11%) or Not Evaluated (14%), underscoring the need for greater taxonomic integration with conservation organizations. Governance advancements in MDD2 include the establishment of external taxonomic subcommittees to guide data collection and curation, a rewritten website that improves access and scalability, a cross-platform application that provides offline access, and new partnerships to continue linking MDD data to global biodiversity infrastructure. By providing up-to-date mammalian taxonomic and nomenclatural data—including links to the text of original name descriptions, type localities, and type specimen collections—the MDD provides an integrative resource for mammalogists and conservationists to more easily track the status of their study organisms.

**Teaser Text:** The Mammal Diversity Database 2.0, listing 6,759 mammal species and 50,230 species-level synonyms, unifies 267 years of taxonomic, nomenclatural, and geographic data to track global mammal biodiversity and provide a continually updated resource for the mammalogical community. **Teaser Image:** Figure 3.

## INTRODUCTION

Species are the central unit of biodiversity research, providing evolutionary and ecological context for zoonotic disease monitoring, conservation policymaking, and other societally critical topics (Cook et al. 2020; Upham et al. 2021; Wang et al. 2021; Colella et al. 2023). Delimiting species is fundamental for understanding biodiversity, allowing researchers to conceptualize meaningful discontinuities among organisms, communities, and ecosystems. Species form the foundation of biodiversity infrastructure that ranges from physical biocollections and digital databases to the knowledge contained in publications (Cook and Light 2019; Galbreath et al. 2019; Heberling et al. 2021). However, species boundaries are constantly changing over time via advances in taxonomy-related fields, leading to the lumping, splitting, and new description of species. In mammals, recent advances in the morphometric, molecular, statistical, and ecological tools used to distinguish previously cryptic taxa, coupled with the increasing globalization of mammalogical research efforts, have led to widespread taxonomic changes (Patterson 2000; Reeder et al. 2007; Burgin et al. 2018; Álvarez-Castañeda et al. 2019).

Although modifying species boundaries to match current evidence of evolutionary relationships is an essential step for studying that history (Sterner 2019), conflict has arisen in mammalogy when various authors have presented different evidence and perspectives on the same taxa (e.g., Frankham et al. 2012; Groves et al. 2017; Zachos 2018a, b; Padial and De la Riva 2021). This plurality of taxonomic views is not unique to mammalogy; instead, this dynamic aspect of biodiversity science emphasizes the importance of rigorous real-time tracking of accepted species names and the associated biological data that signifies their taxonomic meaning (Franz and Sterner 2018; Miralles et al. 2020; Sterner et al. 2020b; Upham et al. 2022a; Sterner et al. 2023). Progress in this arena is critical because identifying which species are recognized, and how they are defined, affects inferred biodiversity patterns at local, regional, and global scales, including assessments of conservation priorities and threats (Mace 2004; Thomson et al. 2018). Thus, broad dissemination of the latest knowledge of species designations and their geographic distributions provides an essential foundation for advancing our collective understanding of mammals globally.

To further this mission, the American Society of Mammalogists launched the *Mammal Diversity Database* (MDD) in 2018, which recognized ∼20% more mammal species (+1,251 and –172 species) compared to the previous authoritative compendium, the *Mammal Species of the World*, third edition (MSW3; Wilson and Reeder 2005; Burgin et al. 2018). Subsequent versions of the MDD have continued to track changes in species- and higher-rank mammal taxonomy as reported in peer-reviewed literature, providing an up-to-date biodiversity resource available online at https://mammaldiversity.org, with past versions archived on Zenodo (MDD 2018, 2019, 2020a, 2020b, 2021a, 2021b 2021c, 2021d, 2021e, 2022a, 2022b, 2022c, 2022d, 2023, 2024a, 2024b, 2024c). The MDD’s rapid online taxonomic updates contrast with the previous reliance on the periodic publication of taxonomic compendia, such as the three MSW editions, nine volumes of the *Handbook of the Mammals of the World* series (HMW), and the two-volume *Illustrated Checklist of the Mammals of the World* (CMW) (Honacki et al. 1982; Wilson and Reeder 1993, 2005; Wilson and Mittermeier 2009, 2011, 2014, 2015, 2018, 2019; Mittermeier et al. 2013; Wilson et al. 2016, 2017; Burgin et al. 2020). Although these compendia provided snapshots of mammalian taxonomy and nomenclature based on published literature and expert opinion, their irregular publication intervals delayed the integration of taxonomic changes and new species descriptions into global biodiversity infrastructure. For example, conservation organizations like the International Union for the Conservation of Nature (IUCN) and community science platforms like iNaturalist rely upon authoritative mammal taxonomies in their decision-making processes (Di Cecco et al. 2021; Cazalis et al. 2022). Establishing the MDD helped create a mammalian taxonomic, nomenclatural, and geographic resource that is comparable to that of other taxon or biome-specific online databases (e.g., World Registry of Marine Species; WoRMS—Vandepitte et al. 2018; Reptile Database; AmphibiaWeb—Uetz et al. 2021; World Flora Online—Borsch et al. 2020; World Checklist of Vascular Plants—Govaerts et al. 2021).

Species’ geographic ranges underlie diverse biological questions, both applied (e.g., where is this species found?) and theoretical (e.g., why are species unevenly distributed?), and are a direct way of communicating the taxonomic meaning of species names. For example, the name *Peromyscus maniculatus* (Wagner, 1845) was long applied to deermouse populations across most of North America, but two proposals in 2019 argued that this species only occurs east of the Mississippi River, with western populations now comprised of *P. sonoriensis* (Le Conte, 1853), *P. labecula* Elliot, 1903, and *P. gambelii* (Baird, 1857) (Bradley et al. 2019; Greenbaum et al. 2019). Because no consistent morphological characters are known to differentiate eastern *P. maniculatus sensu stricto* from the three newly recognized western species, descriptions of each species’ geographic range are essential for communicating this new taxonomic perspective (Sterner et al. 2020a). In general, geographic ranges can be communicated categorically (often via political boundaries; e.g., “Panhandle of Alaska and across N Canada” as used in MSW editions) or using range-map drawings (i.e., geospatial polygons, as used in HMW, CMW, and other compendia). Both types of range information coarsely approximate a species’ true distribution and thus are only accurate at coarse spatial scales, usually ≥100-km resolution (Hurlbert and Jetz 2007; see also recent exchange on this topic by Arbogast and Kerhoulas 2024 and Marsh et al. 2024. In 2008, the IUCN completed the landmark Global Mammal Assessment (GMA), which made range maps available for ∼95% of the species recognized in MSW3 (5,489 species; Schipper et al. 2008). The IUCN Red List subsequently provided the sole source of digital range maps for mammals until Marsh et al. (2022) digitized range maps aligned to three alternative taxonomies: HMW (6,253 species), CMW (6,431 species), and MDD version 1.2 (6,362 species; note that each taxonomy also contains unmapped species that are extinct or data deficient—see Tables 1 and 2 for more detailed totals). Comparison among these datasets documents how the geographic concepts of species-rank mammal taxa have changed over relatively short time periods—mostly from taxonomic changes, in contrast to actual range shifts over longer periods (but see Pacifici et al. 2020). Because geographic knowledge is expected to keep evolving, the MDD taxonomy now tracks categorical changes in the country, continent, and biogeographic realm distributions of accepted species as a core aspect of taxonomic curation.

**Table 1.** Taxon counts for major mammalian taxonomic compendia since 1980 (see Fig. 1 for graphical representation of species totals not including IUCN totals), including the three editions of Corbet and Hill’s (C&H) *A World List of Mammalian Species*, editions four through six of *Walker’s Mammals of the World* (WMW), Anderson and Jones’ *Orders and Families of Recent Mammals of the World* (A&J), Duff and Lawson’s *Mammals of the World: A Checklist* (D&L), three editions of the *Mammal Species of the World* (MSW), *Illustrated Checklist of the Mammals of the World* (CMW), International Union for the Conservation of Nature (IUCN) 2008 and 2024-1 listings, and two major versions of the Mammal Diversity Database (MDD1 and MDD2).

| Taxa | C&H1<br>1980 | MSW1<br>1982 | WMW4<br>1983 | A&J<br>1984 | C&H2<br>1986 | C&H3<br>1991 | WMW5<br>1991 | MSW2<br>1993 | WMW6<br>1999 | D&L<br>2004 | MSW3<br>2005 | IUCN<br>2008<sup>c</sup> | MDD1<br>2018 | CMW<br>2020 | IUCN<br>2024 | MDD2<br>2024 |
| --- | --- | --- | --- | --- | --- | --- | --- | --- | --- | --- | --- | --- | --- | --- | --- | --- |
| **Species** | 4,005<sup>a</sup> | 4,170 | 4,151 | 4,199 | 4,231 | 4,336 | 4,444 | 4,631<sup>b</sup> | 4,809 | 5,115 | 5,416 | 5,489 | 6,495 | 6,554 | 5,983 | 6,753 |
| **Wild Extant** | NA | NA | NA | NA | NA | NA | NA | NA | NA | NA | 5,338 | 5,410<sup>d</sup> | 6,382 | 6,451 | 5,899<sup>d</sup> | 6,629 |
| **Recently Extinct** | NA | NA | NA | NA | NA | NA | NA | NA | NA | NA | 75 | 79 | 96 | 103 | 84 | 113 |
| **Domestic** | NA | NA | NA | NA | NA | NA | NA | NA | NA | NA | 3 | 0<sup>e</sup> | 17 | 20 | 0<sup>e</sup> | 17 |
| **Genera** | 1,014 | 1,033 | 1,017 | 1,057 | 1,055 | 1,066 | 1,116 | 1,135 | 1,192 | 1,117 | 1,230 | 1,241 | 1,316 | 1,343 | 1,308 | 1,353 |
| **Families** | 129 | 135 | 133 | 131 | 132 | 131 | 135 | 132 | 146 | 128 | 153 | 156 | 166 | 167 | 164 | 167 |
| **Orders** | 21 | 20 | 20 | 21 | 20 | 21 | 21 | 26 | 28 | 27 | 29 | 27 | 27 | 27 | 27 | 27 |

<sup>a</sup> Lists 4,007 species, but we identified two duplicates (what is now *Brucepattersonius iheringi* is listed under both *Microxus* and *Oxymycterus*, and *Prosciurillus leucomus* is listed under both *Prosciurillus* and *Callosciurus*)
<sup>b</sup> Corrected total per Solari and Baker (2007)
<sup>c</sup> IUCN 2008 totals based on Schipper et al. (2008)
<sup>d</sup> Includes species classified as Extinct in the Wild and does not include humans
<sup>e</sup> The IUCN does not assess domestic species

**Table 2.** Taxon counts for all 18 versions of the MDD from the initial release of MDD1 in 2018 to the present release of MDD2. Species-level differences (‘diffs’) between versions are noted with the terms ‘de novo’ (additions that coined a new species epithet), ‘split’ (additions that applied an existing species epithet to a newly recognized taxon), and ‘lump’ (removals that combined two or more existing species-rank taxa). Totals of split species do not include names erroneously excluded from earlier versions, and total lumped species do not include names removed due to being considered *nomina dubia* or unavailable. See Supplementary Data SD4 for all MDD version ‘diff’ files and Supplementary Data SD5 for the ‘diff’ file between MDD1 and MDD2 specifically.

| Taxa | MDD1.0<br>Feb 2018 | MDD1.1<br>Mar 2019 | MDD1.2<br>Sep 2020 | MDD1.3<br>Dec 2020 | MDD1.3.1<br>Jan 2021 | MDD1.4<br>Apr 2021 | MDD1.5<br>Jun 2021 | MDD1.6<br>Aug 2021 | MDD1.7<br>Nov 2021 |
| --- | --- | --- | --- | --- | --- | --- | --- | --- | --- |
| **Species** | 6,495 | 6,527 | 6,485 | 6,513 | 6,513 | 6,533 | 6,554 | 6,557 | 6,567 |
| *Wild Extant* | 6,382 | 6,410 | 6,363 | 6,391 | 6,391 | 6,411 | 6,432 | 6,435 | 6,447 |
| *Recently Extinct* | 96 | 100 | 103 | 103 | 103 | 103 | 103 | 103 | 101 |
| *Domestic* | 17 | 17 | 19 | 19 | 19 | 19 | 19 | 19 | 19 |
| *Diff: De Novo* | NA | +22 | +98 | +14 | 0 | +13 | +11 | +5 | +13 |
| *Diff: Split* | NA | +33 | +154 | +15 | 0 | +18 | +10 | +3 | +6 |
| *Diff: Lump* | NA | -31 | -285<sup>a</sup> | -4 | 0 | -10 | 0 | -5 | -8 |
| **Genera** | 1,316 | 1,323 | 1,331 | 1,330 | 1,330 | 1,332 | 1,335 | 1,341 | 1,343 |
| **Families** | 166 | 168 | 167 | 167 | 167 | 167 | 167 | 167 | 167 |

| Taxa | MDD1.8<br>Feb 2022 | MDD1.9<br>Apr 2022 | MDD1.9.1<br>Jun 2022 | MDD1.10<br>Dec 2022 | MDD1.11<br>Apr 2023 | MDD1.12<br>Jan 2024 | MDD1.12.1<br>Jan 2024 | MDD1.13<br>Jul 2024 | MDD2.0<br>Aug 2024 |
| --- | --- | --- | --- | --- | --- | --- | --- | --- | --- |
| **Species** | 6,591 | 6,596 | 6,596 | 6,615 | 6,649 | 6,718 | 6,718 | 6,753 | 6759 |
| *Wild Extant* | 6,471 | 6,476 | 6,476 | 6,494 | 6,526 | 6,594 | 6,594 | 6,623 | 6629 |
| *Recently Extinct* | 101 | 101 | 101 | 101 | 105 | 107 | 107 | 113 | 113 |
| *Domestic* | 19 | 19 | 19 | 20 | 18 | 17 | 17 | 17 | 17 |
| *Diff: De Novo* | +21 | +8 | 0 | +22 | +15 | +37 | 0 | +24 | +2 (296)<sup>b</sup> |
| *Diff: Split* | +6 | +7 | 0 | +26 | +49 | +36 | 0 | +25 | +4 (321)<sup>b</sup> |
| *Diff: Lump* | -3 | -10 | 0 | -30 | -29 | -8 | 0 | -12 | 0 (357)<sup>b</sup> |
| **Genera** | 1,342 | 1,342 | 1,342 | 1,347 | 1,345 | 1,351 | 1,351 | 1,353 | 1353 |
| **Families** | 167 | 167 | 167 | 167 | 167 | 167 | 167 | 167 | 167 |

<sup>a</sup> Most of the ungulate revisions from Groves & Grubb (2011) were removed except where corroborated by later publications
<sup>b</sup> Total differences between MDD1 and MDD2 in parentheses

The MDD’s taxonomic curation also hinges on tracking associated nomenclatural data from historical publications. Nomenclature and taxonomy intersect at the type specimen, which bears the name of the taxon under study (Pyle and Michel 2008). Type specimens provide a stable baseline for subsequent taxonomic revisions given that species names are always linked to name-bearing type specimens (see Definitions Box). The requirement to designate type specimens when defining scientific names, which became common practice in the late 19th century, has created consistency for biological nomenclature. For all animal taxa, the rules of nomenclature are governed by the International Commission on Zoological Nomenclature (ICZN), which created the ICZN Code (ICZN 1999) to standardize how zoological taxonomists create and designate scientific names. Despite historical changes in how researchers define species and other taxa, the system initially proposed by Linnaeus (1758) and now specified by the ICZN Code has allowed for the consistent use of scientific names across centuries of publications. However, despite the critical role of nomenclature in taxonomic curation, it is often difficult for researchers to access these data, including species synonyms, named subspecies, name authority citations, type localities, type specimen identifier(s) and physical location(s) (typically in natural history collections), and original description texts. This is especially true for scientific names that were described in older literature (e.g., before 1900), in journals that have no online repository, in languages other than English, or using type specimens in museums without online catalogs—collectively constituting a type of biodiversity ‘dark data’ whose absence hinders integrative research efforts (Heidorn 2008; Upham et al. 2021). These nomenclatural details are crucial for conducting taxonomic revisions, because identifying whether there is an existing available specific epithet for a potentially valid new species or whether a new name is needed is a key step in the process of defining taxonomic units. The MDD, therefore, jointly tracks the associated taxonomic, nomenclatural, and geographic data that pertain to all mammal scientific names, aiming to make this often difficult-to-access information more accessible and thereby accelerate the process by which integrative taxonomic research can be conducted globally.

In this paper, we introduce version 2.0 of the MDD, a major release that for the first time includes a curated list of species-rank synonyms (including name combinations) for global mammals. We also summarize changes to the public interface and contents of the taxonomic listing during the seven years since MDD version 1.0 was released. We analyze trends that emerged from three of the MDD’s new features: (i) species distributional data, (ii) species-rank synonyms through time, and (iii) associated nomenclatural metadata. We also describe recent collaborations that have greatly improved the quality of MDD data involving the taxonomic and nomenclatural databases the Hesperomys Project (hereafter ‘Hesperomys’; https://hesperomys.com) and Bats of the World (hereafter ‘Batnames’; https://batnames.org). These collaborations highlight the importance of data sharing and the need to continue building open forums for communicating biodiversity information globally, which can dramatically streamline taxonomic research efforts. Lastly, we describe recent and upcoming changes to the MDD governance structure to invite greater editorial oversight from globally distributed groups of taxonomic experts. The MDD aims to continue providing an authoritative, open-access, and community-driven resource for advancing mammalogical research and conservation worldwide.

## METHODS

Version 1.0 of the MDD (MDD1) was released in 2018 with a taxonomic cutoff date of 15 August 2017 (Burgin et al. 2018). Support from the American Society of Mammalogists (ASM) and VertLife Terrestrial NSF project (DEB: 1441737) enabled the creation of an online database with the aim of providing a continuously updated compendium of the world’s extant and recently extinct mammal species and higher-rank taxa. Since then, the MDD database and public web interface have been continually hosted at https://mammaldiversity.org with species-specific pages that include comments and citations relating to taxonomic and nomenclatural changes that have occurred since the MSW3 cutoff of 2004 (Wilson and Reeder 2005). Additional content of the MDD1 release included valid species names, authority author and year, species higher-rank taxonomy, biogeographic realm distribution, extinct/extant status, and taxonomy notes. Subsequent versions of the MDD added species IUCN conservation statuses, English common names, and type localities (version 1.2). Coarse synonym listings also began being curated with version 1.2, consisting of all unique available and unavailable species-rank names included under each species and their respective authorities (previously called ‘nominalNames’), as did type specimens and geographic categories of country and offshore territory distributions. The present publication describes the version 2.0 release of the MDD (MDD2), which focuses on summarizing the global patterns that arise from the more recent fields added to the MDD, including data pertaining to nomenclatural availability, type localities, type specimens, original description citations and links, and original name combinations when available for each species and synonym. A cutoff date of 15 August 2024 was set for including taxonomic, nomenclatural, and geographic distribution changes in MDD2.

During the seven years since MDD1 was released, 16 incremental releases of the MDD (Table 2) have been published as CSV files with associated metadata and are archived on Zenodo (see Supplemental Data SD3 for complete species lists of all previous MDD versions). Since MDD version 1.3.1, a separate ‘diff’ CSV file has listed major changes between the current and previous versions. Starting with MDD2, a synonym-specific CSV file is included that lists every species-rank scientific name applicable to extant and recently extinct mammal species, including all associated nomenclatural data for each name. Collaborations with other biodiversity databases have been critical for expanding and improving the curation of data included in the MDD. Integration of nomenclatural data from Hesperomys (Zijlstra 2024) – an online database cataloging the nomenclature of living and extinct animals, with a focus on mammals – has greatly strengthened the MDD. These data were primarily integrated into the MDD species-rank synonym data, resulting in expanded coverage of type localities, type specimens, and other nomenclatural data detailed below. Coordination with Batnames (Simmons and Cirranello 2024) has also been critical to standardize the taxonomic arrangement of the Order Chiroptera so that, starting in January 2024 with MDD v1.12 and Batnames v1.5, all MDD species- and higher-rank taxa of bats are matching between these databases (however, subspecies of bats are currently only listed in Batnames). A further collaboration with Yale University researchers enabled the digitization of geographic range maps as polygon shapefiles for all mammal species, as matched to the taxonomies of MDD v1.2, HMW series, and CMW (see Marsh et al. 2022 and map downloads updated to MDD v1.4 via the ‘mddmaps’ R package; Robles-Fernández 2024). Lastly, efforts to coordinate with the ASM Mammal Images Library have aligned the taxonomic names and concepts used in the expert-identified images to each new release of the MDD (see https://www.mammalogy.org/image-library).

The MDD’s taxonomic, nomenclatural, and geographic data will continue to be updated with new publications in forthcoming database releases. Continued changes are expected regarding species’ taxonomic status, nomenclatural information, geographic distributions, and common names, given the steady rate of new taxonomic evidence being gathered and integrated relative to older publications. The MDD is a living database, and feedback from mammalogists regarding errors, missing information, and potential updates is openly welcome. The MDD currently accepts public feedback in two ways: (i) emails to; and (ii) GitHub issues: https://github.com/mammaldiversity/mammaldiversity.github.io/issues. Feedback is reviewed and acted upon regularly by the MDD team.

### Taxonomic Decision-Making

A fundamental goal of the MDD has been to maintain an updated taxonomic checklist of the world’s currently recognized extant and recently extinct mammal species. To do so, the MDD team has continuously tracked mammal taxonomic literature with the aid of Google Scholar email alerts, periodic examination of journal webpages, and users directly sending publications. The MDD curation team, including all authors of this paper, consists of student research assistants and web developers (paid by the ASM), along with volunteers and consultants. This group is collectively advised by the ASM Biodiversity Committee (https://www.mammalsociety.org/committees/biodiversity). MDD taxonomic curation has emphasized changes since the publication of MSW3 in 2005. Most taxonomic changes incorporated into the MDD have, thus far, closely followed the most recent empirical evidence given in peer-reviewed articles, which each represent the current best hypothesis of species boundaries. By consistently following the most recent evidence from peer-reviewed literature, the MDD’s taxonomy is designed to collectively reflect the most up-to-date taxonomic perspective of the global mammalogical community and thereby minimize subjective biases coming from the MDD team.

Nevertheless, occasionally controversial issues in the peer-reviewed literature have required the MDD team to make subjective decisions regarding the most justified current taxonomy. In cases where there is insufficient evidence to support a taxonomic change or if multiple conflicting taxonomic arrangements are published in rapid succession, the MDD team provides justifications and citations on the relevant per-species pages with summaries given on the About page at https://mammaldiversity.org. In cases of ongoing controversy, a ‘flag’ designation is noted in the CSV file. Most subjective taxonomic decisions have been made internally by the MDD team. However, beginning with v1.10 (December 2022), the MDD piloted a protocol for publishing taxonomic opinion ‘white papers’ on Zenodo in collaboration with members of the Global Bat Taxonomy Working Group, which is part of the IUCN SSC Bat Specialist Group and aligned with Batnames. Three subjective decisions were issued: (i) on the species status of *Myotis evotis* in relation to *Myotis keenii* (Lausen et al. 2019, 2021; Morales et al. 2021) by Upham et al. (2022b), (ii) on recognizing *Aeorestes*, *Dasypterus*, and *Lasiurus* as either subgenera or genera (Baird et al. 2015, 2017, 2021; Ziegler et al. 2016; Novaes et al. 2018; Teta 2019) by Francis et al. (2023), and (iii) on whether the five *Myotis lucifugus* subspecies should be recognized as subspecies or species (Morales and Carstens 2018) by Francis et al. (2022). This third white paper was issued without MDD involvement but is still followed in the MDD taxonomy. The goal of formally issuing subjective decisions is to transparently evaluate the available taxonomic evidence, present a detailed justification for choosing one taxonomic arrangement over another, and then make recommendations for gathering evidence to reach more definitive conclusions. The approach of self-publishing on Zenodo was chosen to avoid furthering arguments in the peer-reviewed literature and to incentivize stakeholders to gather new data to support their claims.

Two other major subjective decisions have been made by the MDD team. The first was to exclude almost all the new taxonomic arrangements proposed in Groves and Grubb’s *Ungulate Taxonomy* (Groves and Grubb 2011; Groves et al. 2011; Burgin et al. 2020—change made in MDD v1.2). Groves and Grubb (2011) recommended a large number of taxonomic changes within Perissodactyla and terrestrial Artiodactyla that were largely based on qualitative morphological characteristics with small sample sizes. These changes have been controversial within the mammalogical community, and many specialists have subsequently reverted to the taxonomy presented in MSW3 (e.g., Holbrook 2013; Gutiérrez and Garbino 2018). The exception was for the MDD to retain changes that have been subsequently validated by peer-reviewed research (e.g., *Nanger* gazelles; Lorenzen et al. 2007; Siegismund et al. 2013).

Second, the MDD excluded several recently suggested taxonomic changes and new species, subspecies, and generic names described by Raymond Hoser, who has prolifically named species in a self-published journal, generally using data from other publications as justification. These changes were previously restricted to reptiles, amphibians, and a few spiders and fish, but 46 mammalian names were described between 2018 and the MDD2 cutoff date. Based on the suggestions of Wüster et al. (2021), the MDD considers these names as examples of ‘taxonomic vandalism’ that should be ignored other than being listed in the synonym dataset (given the nomenclatural status of ‘rejected_by_fiat’ to signify that the broader taxonomic community rejects these names, not the ICZN Code).

### Species Pages and Distributional Data

Each species in the MDD has a separate webpage that includes the accepted scientific name; authority author(s) and year; higher-rank taxonomic arrangement; primary and alternative English common names; original description citation and link to original description article; type locality, type specimen catalog number, and link to the type specimen in a natural history collection; list of species-rank synonyms and associated nomenclatural data; country-level distribution and accompanying map of listed countries; taxonomic notes and associated citations; IUCN Red List status (see *Conservation Status* section below); whether the species is extant or extinct, wild or domesticated, or has a ‘flagged’ designation denoting an ongoing taxonomic controversy; and an associated permanent hyperlink (permalink) to access the species page directly. The nomenclatural data presented on species pages are sourced from the synonym CSV file, and those data are detailed under the *Nomenclatural Metadata* section below.

#### Higher-rank Taxa

The MDD also tracks the higher-rank taxonomy of living and recently extinct mammal species, including the ranks of class, subclass, infraclass, magnorder, superorder, order, suborder, infraorder, parvorder, superfamily, family, subfamily, tribe, genus, and subgenus. All currently recognized species on the MDD have been assigned within an order, family, and genus, with intermediary higher and lower ranks assigned when suggested in the literature. For now, taxonomic changes at the family and genus ranks are noted in the taxonomic notes for species affected by those changes, while changes at the order rank are noted in the ‘About’ page on the MDD website (e.g., use of the order names Eulipotyphla and Artiodactyla instead of Lipotyphla and Cetartiodactyla; Asher and Helgen 2010). In cases where an intermediate rank is used (e.g., subfamily, tribe) and there are lower-rank taxa that are not confidently placed within that intermediate rank, the term *incertae sedis* is used. For example, within the subfamily Sigmodontinae, most genera are grouped within tribes, but the genus *Abrawayaomys* has yet to be placed into any recognized sigmodontine tribe (Ventura et al. 2013), so the tribal affiliation for that genus is *incertae sedis*.

#### Common Names

The primary English common names (also called vernacular names) for all species were initially sourced from the HMW and CMW series, which employed a standardized naming convention. Because common names are not formally regulated for mammals, there is some leeway for modifications. However, the MDD aims to maintain consistent naming conventions. The primary common name reflects the most widely used or uncontroversial common name for a species while avoiding repeated common names or ‘names-within-names’ (e.g., ‘Common Hippopotamus’ rather than ‘Hippopotamus’ used for *Hippopotamus amphibius* to avoid potential confusion with the ‘Pygmy Hippopotamus’, which also includes the term ‘Hippopotamus’). For species not included in the HMW or CMW series or that required a common name change due to taxonomic revisions, the MDD team generates a new or revised common name (e.g., when the Nine-banded Armadillo, *Dasypus novemcinctus*, was split into multiple species, the MDD used ‘Mexican Long-nosed Armadillo’ as the common name for *Dasypus mexicanus* and ‘Southern Long-nosed Armadillo’ for *Dasypus novemcinctus sensu stricto*; Barthe et al. 2024). Common names suggested by the species name authority or taxonomic change publications are prioritized, but if no common name has been suggested or if the suggested name conflicts with the primary common name of another accepted species, then a new common name based on the species’ diagnostic features, geographic distribution, or scientific name etymology is generated. The MDD also maintains consistent common name components within each genus when appropriate (e.g., ‘deermouse’ for all *Peromyscus* species) and capitalizes all words of the common name as proper nouns (e.g., Aztec Deermouse for *Peromyscus aztecus*). The ‘other common names’ field includes other known common names that have been used in English either widely or locally or used for subspecies within that species, as initially sourced from the HMW and CMW series, with additional names added as they were found while searching the literature. The MDD also lists names from other languages if they are widely used, but this list is not exhaustive. The MDD team expects to continue adding and revising common names regularly as they are found or updated.

#### Conservation Status

IUCN Red List statuses (Least Concern [LC], Near Threatened [NT], Vulnerable [VU], Endangered [EN], Critically Endangered [CR], Extinct in the Wild [EW], Extinct [EX], Data Deficient [DD], Not Evaluated [NE]) are listed for each species, as currently sourced from version 2024-1 of the IUCN Red List (IUCN 2024a). If a species is recognized under a different scientific name by the IUCN than what is used in the MDD (e.g., different genus or spelling differences), the scientific name used by the IUCN is given in parentheses following the status in the ‘iucnStatus’ field (e.g., ‘VU (as *Onychogalea fraenata*)’ for *Onychogalea frenata*).

Additional fields with binary coding are included for whether a species is extant or recently extinct and wild or domesticated. All species that are believed to have been extant after the year 1500 CE according to evidence published in the peer-reviewed literature are included in the MDD. The representatives of 17 mammal species that were domesticated by humans during prehistoric times are considered separate species from their ancestors in the MDD, all but two of which (*Bos primigenius* and the unknown ancestor of *Camelus dromedarius*) are extant. The MDD’s recognition of these domestic forms as distinct species contradicts much of the evolutionary and taxonomic research on these taxa, which has shown that the wild and domestic forms should be considered a single species in most cases (e.g., Driscoll et al. 2007; Groenen 2016; Jackson et al. 2019, 2020). However, the MDD has chosen to recognize these specific domestic forms as distinct species to avoid confusion regarding which forms are being discussed in both political and conservation situations. Recognizing domestic species also emphasizes the wild forms as an entity to be separately considered and protected. Thus, the MDD acknowledges that recognizing these 17 domestic forms as species is a deviation from the species recognition standards outlined in the *Taxonomic Decision-Making* section above and follows the naming conventions for domestic species and their wild ancestors outlined by Gentry et al. (2004).

#### Geographic Distributions

The MDD started tracking the native continental and country and offshore territory distributions of all currently valid and wild mammal species starting with v1.2. As of MDD2, four scales of geographic distribution are documented: (i) countries and offshore territories, (ii) country subregions, (iii) continents, and (iv) biogeographic realms. The regional distributions of all terrestrial species are defined by individual organisms physically existing within the boundaries of the defined region. Aquatic marine or freshwater species (all cetaceans, sirenians, pinnipeds, and marine mustelids) are defined as existing either along the shoreline of a given region or in an inland water body within or bordering the defined region; the one exception is for biogeographic realm distributions, where aquatic species are only included if they are found in inland water bodies within the biogeographic realm (marine distributed species are listed as ‘Marine’ in that distribution field).

Species distributional data was initially sourced from the IUCN, HMW, and CMW, as supplemented with records published in peer-reviewed literature and regional taxonomic and geographic compendia. Recently introduced and extinct populations are not included in the distribution of living mammals, although the former distribution is listed for globally extinct species, as well as ancient introductions (before 1500 CE) of extant species (e.g., Iberian populations of *Genetta genetta* represent ancient human-introduced populations and are included in the MDD; Delibes et al. 2019). All distributional listings are generally supported by a confirmed record of the species living there (i.e., a museum specimen or photographic evidence), although, if a species is suspected to occur in a region but there is no evidence to confirm its presence, a question mark is listed after the region name. A coarse country- and offshore territory-level map is also included on the webpage for each species—this should not be confused with a more fine-grained ‘expert range map’ of the species, which are currently not hosted directly by the MDD (but see the R package by Robles-Fernández 2024). Region naming conventions generally follow the International Organization for Standardization (ISO), with occasional exceptions to match abbreviated naming conventions or for regions not covered by the ISO (see Supplementary Data SD7 for a list of regions used in the MDD).

The list of countries and offshore territories includes all United Nations member and observer states, as well as relevant offshore territories geographically separated from their governing country (e.g., French Guiana is listed separately from France). Country subregions include all the states, provinces, territories, and other applicable subregions for larger countries. As of MDD2, country subregions of species have only been implemented for the United States (all 50 states and the District of Columbia). However, country subregions have been fully implemented in tracking the higher geography of type localities on the synonym sheet (see *Nomenclatural Metadata* section below).

MDD species distributions are coded for seven continents with the following definitions: North America (North and Central America to the border between Panama and Colombia, Greenland, Bermuda, and all of the Greater and Lesser Antilles except Aruba, Bonaire, Curaçao, and Trinidad and Tobago); South America (continent south of the border between Panama and Colombia, including Aruba, Bonaire, Curaçao, Trinidad and Tobago, Galapagos Islands, and Falkland Islands); Europe (Eurasia west of the Ural mountains and river, north of the Caucasus mountains, and north and west of the Black and Caspian seas, including the Azores and all the islands in the Mediterranean except Cyprus); Asia (Eurasia east of the Ural mountains and river, east of the Bosporus and Dardanelles and including Cyprus, south of the Caucasus mountains, and south and east of the Black and Caspian seas, east of the Suez Canal, and west of Weber’s Line, including Christmas Island); Africa (continental Africa west of the Suez Canal, including Madagascar, Madeira, Canary Islands, Ascension, Tristan da Cunha, and Saint Helen, amongst other offshore islands); Oceania (continental Australia, and Melanesia, Polynesia, and Micronesia east of Weber’s Line, including Hawai’i); and Antarctica (continental Antarctica and all Subantarctic islands, including South Georgia and the South Sandwich Islands).

The MDD considers eight terrestrial biogeographic realms, as defined by Olson et al. (2001): Nearctic, Neotropical, Palearctic, Afrotropical, Indomalayan, Australasian, Oceanian, and Antarctic; because these are terrestrial biogeographic realms, marine distributed species are listed as ‘Marine’. In some cases, the MDD deviates from Olson et al. (2001) when a small portion of a biogeographic realm is geographically separate from the primary area of the realm, such as including southern Florida in the Nearctic rather than Neotropical realm and southern Iran in the Palearctic rather than Afrotropical realm.

### Nomenclatural Metadata

MDD2 includes a newly curated list of species-rank synonyms – created in collaboration with Hesperomys (Zijlstra 2024) – for every mammal species in the MDD, totaling 50,230 synonyms. The full synonym list is available on Zenodo and as Supplementary Data SD2. The synonym list includes available and unavailable names; spelling variants and name combinations; and names that are associated with currently valid species, subspecies, synonyms, *nomina dubia*, or *species inquirendae*. Nomenclatural availability of each name was determined based on application of the ICZN Code (ICZN 1999) as well as executive decisions made by the ICZN on the protection or rejection of specific names when applicable. The MDD follows the ICZN Code to inform issues of the nomenclatural availability and spelling of scientific names, which has led to changes in the spelling of specific epithets on the MDD that occasionally differ from the published literature, typically involving changes in gender agreement. These cases are usually noted in the taxonomic notes for species if these changes affect the spelling of a valid species name, and all such spelling changes between MDD1 and MDD2 are listed in Supplementary Data SD5.

#### Literature Search

Nomenclatural information included in both the MDD and Hesperomys was collected by manually and programmatically searching the primary taxonomic literature, *Mammalian Species* accounts, and secondary global (e.g., MSW, HMW, CMW) and regional taxonomic compendia (e.g., Gardner 2008; Jackson and Groves 2015; Patton et al. 2015).

Baseline synonym information was sourced from MSW3, which listed the specific epithet and authority for the synonyms of each species (Wilson and Reeder 2005). These searches spanned multiple online publication repositories, including Google Scholar (https://scholar.google.com), the Biodiversity Heritage Library (https://www.biodiversitylibrary.org), HathiTrust (https://www.hathitrust.org), the Internet Archive (https://archive.org), and Gallica (https://gallica.bnf.fr).

The primary goal of this ongoing search is to verify the existence, nomenclatural status, and associated data of every name applied to an extant or recently extinct mammal species. In cases where the MDD team has found an earlier citation for a name, differences in spelling or authority, or differing type localities from those listed in recent literature, this information is updated between the MDD and Hesperomys synonym datasets, along with the valid species if applicable. These efforts have also located names that were not included in recent taxonomic compendia (e.g., MSW3), which are added to the MDD-Hesperomys synonym dataset. Each name has been associated with its original description citation, including the original name combination, page of first use, type locality, and type specimen when available. Much of the initial nomenclatural data and citations were sourced from the major compendia mentioned above, but these sources often provided abbreviated citations that can be difficult to decipher and track, which required additional data acquisition directly from the original references.

#### Names and Authorities

The MDD catalogs the verbatim original name combination from the original description of each name, as well as the taxonomic rank the name was initially described as and the current spelling of the specific epithet based on application of the ICZN Code (termed the ‘rootname’). The MDD lists the currently accepted primary higher taxonomy (order, family, genus, and species) from the species data for each name, using *incertae sedis* for names not assignable to any of these taxonomic ranks (i.e., *nomina dubia*, *species inquirendae*). The authority for each scientific name is given in a standardized format, including parentheses around the authority name and year when applicable (see Definitions Box). All authors in the author line are included in the authority and an ‘in’ statement is used when an authority is listed in the publication that differs from the author line (e.g., Kitchener in Kitchener & Maryanto). When multiple authors have the same surname, their first and middle initials are included before their surname to differentiate between each of them (e.g., J. A. Allen versus G. M. Allen; J. Eric Hill versus J. Edwards Hill).

#### Original Citations

To verify each name’s citation and associated nomenclatural data, the MDD team has examined the original descriptions of each name, including the page number where the name or description first occurs and the exact month or day of publication when available. Citations are considered verified when either a PDF of the cited source is obtained or when a member of the MDD team has physically examined the cited source. If the citation has not been verified, the MDD usually lists citation information found in other publications and compendia until the citation can be verified.

When available, the MDD also links names to an online version of the original description publication, as well as the corresponding page on Hesperomys. For most names initially described or mentioned in publications published before 1925, the MDD team searches for links on the Biodiversity Heritage Library (BHL). Because the BHL provides permalinks associated with individual pages of digitized publications, links can be found for the exact page on which the name was first mentioned or described. Many of these links were found programmatically by using metadata in the BHL, such as the original description volume and page numbers (often sourced from compendia) and then were either verified programmatically or manually by checking for the presence of the name in the text for the page on the BHL website. For older publications not available on the BHL but available from other online literature repositories (e.g., HathiTrust, Internet Archive, Gallica), the MDD provides manually sourced direct links to either the first page on which the name was mentioned or to the entire paper when no page-specific links are available. For names described in more recent publications (usually after the year 2000), the MDD provides a Digital Object Identifier (DOI) linking the name to the journal article’s webpage instead of a direct link. No link is provided when an original citation is not available online or has not yet been located by the MDD team, although the citation might have been verified by other means of obtaining a Portable Document Format (PDF) file (e.g., through interlibrary loan).

#### Nomenclatural and Taxonomic Status

For each name, the MDD summarizes both the nomenclatural and taxonomic status. The nomenclatural status field includes an array of designations relating to the ICZN Code, such as whether a name is available or unavailable (and why it is unavailable if applicable) or if it is preoccupied or a *nomen novum*, spelling variant, or name combination (see Definitions Box). For names that are variants of previously published names, such as spelling variants and name combinations, the MDD records the accepted species name for which it is a variant. For names that are preoccupied by an earlier name through primary or secondary homonymy, the MDD records the preoccupying name. The validity field lists whether a name is a valid species name, a synonym of a valid species, currently unassigned to a single valid species, or having type material of composite or hybrid origin. The MDD also plans to include whether a name is a valid subspecies in future releases but has thus far refrained from tracking subspecies designations due to their inconsistent use and multiple definitions in the published literature (see Burbrink et al. 2022). See Supplementary Data SD8 for a list of all nomenclatural and validity statuses in the MDD2 and Hesperomys synonym listing.

#### Type Localities

The MDD records type localities for all applicable names as they are added to the MDD synonym sheet. Type localities were initially sourced from the HMW and CMW series, which listed the localities in a standardized format and included type locality restrictions and citations. On the current synonym sheet, the MDD has now verified and copied the verbatim type locality from the original description text for names with verified citations. When the MDD has not verified the verbatim original type locality, type localities are listed as obtained from other primary and secondary sources.

The MDD team has placed each type locality into its modern country borders (or higher geographical region when no country was specified or determinable from the original description). For many larger countries, the MDD also records the geographic subdivision where the type locality is located (e.g., US states, Chinese provinces, specific islands of archipelagic countries like the Philippines and Indonesia). The region naming conventions and boundaries follow the same conventions as the distributional data of species-rank taxa described in the *Geographic Distributions* section above. In cases where geographic coordinates are given for the type locality in the original description or later publications, the MDD separately records these in decimal degrees. For all names described since the year 2000 that did not already have coordinates present in the original description publication, the MDD team geolocated the type locality using GeoLocate (www.geo-locate.org) or WikiMedia Geohacks location coordinates.

*Type Specimens*.

The MDD and Hesperomys aim to link each available name with its type material. These links are often sourced from type catalogs published by various museums (e.g., Lawrence 1993; Borisenko et al. 2001), along with original species descriptions and taxonomic revisions. The MDD lists type specimens in a standardized format for each institution, along with the kind of type specimens (holotype, lectotype, neotype, syntypes, or nonexistent). When available, the MDD provides a link to the online collection database for the institution where the specimen is housed. In some cases, only the collection where a specimen is located could be determined and not the complete specimen number, in which case it is labeled as ‘unnumbered’.

### Data Presentation and Applications

Several changes have been introduced to the MDD website with the release of MDD2, including a new cross-platform application: the MDD app. Co-author HH wrote the MDD app using the Flutter framework (https://flutter.dev/) and the Rust programming language (https://www.rust-lang.org/). It supports iOS, iPadOS, Android, Windows, Linux, and macOS. Details on installing the app are available at https://github.com/mammaldiversity/mdd_app. Critically, the MDD app allows offline access to MDD data, which makes it convenient to access during remote fieldwork when an internet connection is unavailable. The MDD app also features an export function for a whole or subset of MDD data to a CSV file, which can then be shared. We revamped the website, including a migration from Jekyll (https://jekyllrb.com/) to the Astro web-framework (https://astro.build/) with TypeScript (https://www.typescriptlang.org/), and Tailwind CSS (https://tailwindcss.com/) integration. The new website offers better extensibility to add new features, improved user interface and performance, and the integration of taxonomic synonyms. The website layout is adaptive to the screen size of the browser, allowing for easy viewing on small devices such as smartphones or tablets. This layout also supports dark and light themes that are adaptive to the users’ browser appearance setting. The website provides a permalink for each taxon that third-party websites can easily access given that it follows a standard format using the MDD ID for each taxon. For example, the permalink for *Ornithorhynchus anatinus* is https://www.mammaldiversity.org/taxon/1000001.

### Analyses

To summarize taxonomic trends, the MDD2 taxonomy was compared to other major compendia, including Corbet and Hill’s first (C&H1), second (C&H2), and third (C&H3) editions of *A World List of Mammalian Species* (Corbet and Hill 1980, 1986, 1991), editions four through six of *Walker’s Mammals of the World* (WMW; Nowak and Paradiso 1983; Nowak 1991, 1999), Anderson and Jones’ *Orders and Families of Recent Mammals of the World* (A&J; Anderson and Jones 1984), Duff and Lawson’s *Mammals of the World: A Checklist* (D&L; Duff and Lawson 2004), MSW1 (Honacki et al. 1982), MSW2 (Wilson and Reeder 1993), MSW3 (Wilson and Reeder 2005), CMW (Burgin et al. 2020) and the IUCN (IUCN 2024a), as well as across all 18 versions of the MDD. Additionally, total species counts were extracted from each source (other than IUCN version totals, which are not representative of all recognized mammals at their time of versioning) and used to estimate the rate of new mammal species recognition from 1980 to 2024 using linear regression. The equation generated for the linear regression line was then used to estimate total recognized mammal species by 2030, 2050, and 2100 if the observed trends were to continue. Corbet and Hill (1980) is the first publication to include a complete list of contemporarily recognized mammal species in the 20th century, which is why we use that publication as the starting point.

The dates of all available name and valid name descriptions were plotted to visualize the history of species-rank mammal descriptions since 1758. The proportion of species-rank names that were originally described as either species or subspecies and names that have been either lumped or split since their original descriptions were also summarized using 10-year running means from 1 January 1758 to 15 August 2024. Geographic trends in accepted species richness, conservation status (threatened and understudied species), species endemism, new species description, and available name type localities were also evaluated, including across continents and biogeographic realms. State-level species richness across the United States and the global distribution of type localities for names described after the year 2000 were plotted to investigate biodiversity patterns. Lastly, total species per IUCN Red List category were summarized to estimate the proportion that are Low Risk (LC and NT species), Understudied (DD and NE species), Threatened (VU, EN, CR, EW species), and Extinct (EX species) according to the IUCN. The R code used to analyze these data, plot figures, and create supplementary files is available in a GitHub repository accessible at https://github.com/mammaldiversity/MDDv2_ms.

## RESULTS

### Taxonomic Change and Global Biodiversity

MDD2 catalogs the taxonomy and distribution of 6,759 valid mammal species (6,629 wild extant; 113 extinct; 17 domestic), along with nomenclatural data for 50,230 valid and synonymous species-rank scientific names (Fig. 1; summarized in the *Nomenclatural Metadata* section below). MDD2 includes 267 more species (+4.1% over 7 years) than the 6,495 species recognized in MDD1 and 1,343 more (+24.8% over 20 years) than the 5,416 in MSW3 (Table 1). In contrast, the 2024-1 IUCN Red List includes 5,983 species, which is 776 fewer (−11.5%) than MDD2 despite being released the same year. The ∼25% increase over 20 years in recognized mammal species is comparable to the 31.1% increase (+1,246 species) over a 22-year period from the 1982 release of MSW1 to the 2004 taxonomic cutoff of MSW3. Over a broader time period, MDD2 recognizes 2,754 more species (+68.8% over 44 years; 3,149 splits + de novo, 395 lumps) than the 4,005 species recognized in 1980 by C&H1, which was the first modern mammal taxonomic compendium to include a complete list of living and recently extinct mammal species. Linear regression of estimates across 14 compendia recovered a significant increase in mammal species recognition from 1980 to 2024 (y = 64.86x – 124581.37; R^2^ = 0.98; Fig. 1). Projecting this rate of species recognition out to the future, there will be ∼7,084 species by 2030, ∼7,733 species by 2040, ∼8,382 species by 2050, and ∼11,625 species by 2100.

**Figure 1.**
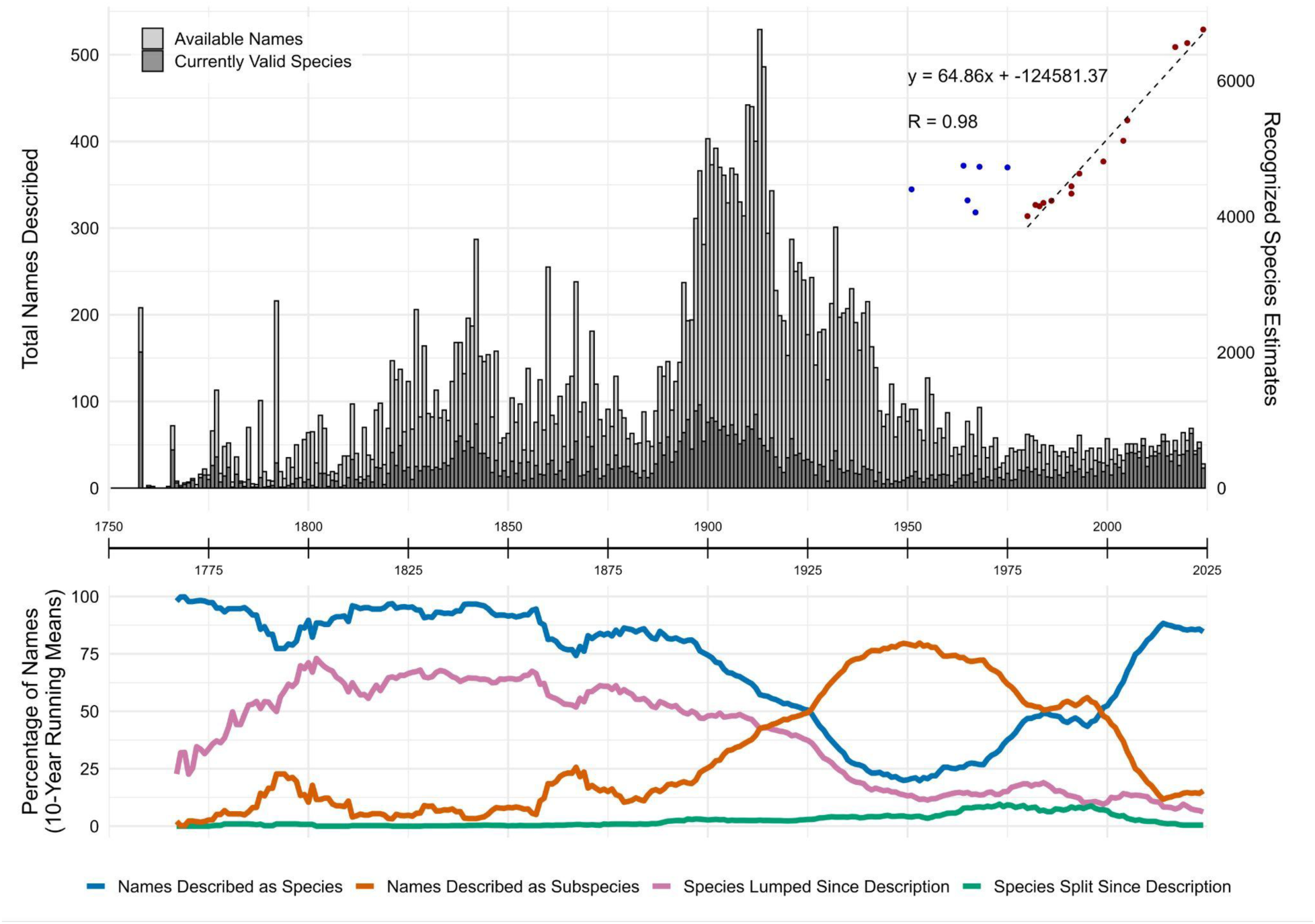
A graphical history of mammalian species-group name descriptions since the start of zoological nomenclature on 1 January 1758, incorporating all 28,382 available names (including preoccupied, replacement, and suppressed names) and 6,759 names currently recognized as valid species in MDD2. The histogram summarizes the number of available names (light gray) and valid species names (dark gray) described each year from 1 Jan 1758 to 15 Aug 2024 (a ∼267 year period). The lines in the bottom panel summarize the 10-year running means (starting 1768) for the proportion of names initially described at the ranks of species (blue) and subspecies (orange), as well as the proportions of names initially described as species and then lumped into another species (pink) or initially described as subspecies, forms, or varieties and then split into their own species (turquoise). Since all four lines were smoothed by taking 10-year running means, each point on these lines represents a mean generated from the previous 10 years of data. See Supplementary Data SD9 for the raw data and running means used to create the line graph in this figure. Total recognized mammal species estimates are included from major compendia since Corbet and Hill (1980; red points), including all taxonomic compendia listed in Table 1 other than IUCN totals (which are not representative of the actual recognized mammals at the time of versioning). A linear regression line, equation, and R^2^ value of these data are included on the graph. Six additional pre-1980s species estimates are also included for comparison (blue points), which are from resources that do not include full species lists (4,400 species in Storer 1951; 4,748, 4,732, and 4722 species in Walker et al. 1964, 1968, 1975; 4,237 species in Morris 1965; 4,060 species in Anderson and Jones 1967).

Details of the full MDD2 taxonomy, including associated citations, metadata, and geographic assignments, are provided in Supplementary Data SD1. Additionally, a count-based summary of taxon numbers, available names, new species, and IUCN status categories per mammal family and order in MDD2 is provided in Supplementary Data SD6.

The 18 total releases of the MDD from MDD1 to MDD2 included 15 primary releases (i.e., 1.X and 1.XX releases) and 3 patch releases (i.e., 1.X.X releases) from Aug 2017 to Aug 2024 (Table 2). Over this period, the MDD cataloged a total of 305 de novo species (newly described names), 392 split species (existing synonymous or subspecies names reevaluated as valid species), and 435 lumped species (species subsumed into synonymy at the species rank), which includes some taxa that were first split and re-lumped and vice versa across versions (see Table 2 for taxon totals and version differences, and Supplementary Data SD4 for all per-version MDD ‘diff’ files). When comparing the differences between MDD1 and MDD2 directly (see Supplementary Data SD5 for an MDD1-to-MDD2-specific ‘diff’ file), there were 296 de novo species, 321 split species, 357 lumped species, 11 missing species that were subsequently added (mostly recently extinct and domestic species), 6 removed species (all either extinct species that did not survive recently enough for inclusion, or names now considered unavailable or *nomina dubia*), 227 generic changes, 17 specific epithet name changes, and 148 spelling changes and corrections. The highest number of taxonomic changes between two MDD versions occurred between versions 1.1 (Mar 2019) and 1.2 (Sep 2020), versions 1.9.1 (Jun 2022) and 1.10 (Dec 2022), and versions 1.10 and 1.11 (Apr 2023) (Table 2). The largest single-version difference, between versions 1.1 and 1.2, is largely attributed to the removal of the Groves and Grubb (2011) taxonomy for Perissodactyla and terrestrial Artiodactyla (see *Taxonomic Decision-Making* section in Methods), which makes MDD v1.2 the only version that cataloged more species lumps than additions. Version 1.11 is also notable because it marked the beginning of collaborations between the MDD and Hesperomys and Batnames, which involved considerable efforts to reconcile information across the three databases, leading to many changes in taxonomy (15 de novo; 49 splits; 29 lumps) and nomenclature (e.g., spelling changes to 52 species-rank specific epithets) despite a gap of only 5 months between MDD v1.10 and v1.11 release dates.

Between MDD1 and MDD2, there were net differences of 37 more genera (1,316 to 1,353) and 1 more family (166 to 167) recognized, with no changes to the 27 orders. The families Callitrichidae and Aotidae were recognized as distinct, while Eschrichtiidae was lumped into Balaenopteridae since MDD1. In contrast, the MDD2 taxonomy documents a net increase of 339 (27.6%) genera, 38 (29.5%) families, and 6 (28.6%) orders since C&H1 and a net increase of 123 (10.0%) genera and 14 (8.4%) families since MSW3 (Table 1). MDD2 recognizes 2 fewer orders than MSW3 because Soricomorpha and Erinaceomorpha are now lumped into a single order (Eulipotyphla), and Cetacea is now lumped into Artiodactyla. The IUCN version 2024-1 recognizes 1,308 genera and 164 families, which is a net difference of 45 (3.3%) fewer genera and 3 (1.8%) fewer families than MDD2 but recognizes the same 27 orders. MDD2 includes 8 families that the IUCN does not (†Archaeolemuridae, Cetotheriidae, Choloepodidae, Heterocephalidae, †Megaladapidae, Potamogalidae, Zapodidae, and Zenkerellidae) while the IUCN includes 5 families that MDD2 does not (Capromyidae, Eschrichtiidae, Megalonychidae, Myocastoridae, Neobalaenidae—note that Cetotheriidae/Neobalaenidae and Choloepodidae/Megalonychidae are alternative names for the same extant species compositions; Fordyce and Marx 2013; Marx and Fordyce 2016; Delsuc et al. 2019). See Table 1 and 2 for further generic, familial, and ordinal comparisons among major mammal compendia published since 1980 and among MDD versions.

The three most speciose mammal orders are still Rodentia (2,747 species; 536 genera; 35 families), Chiroptera (1,485 species; 236 genera; 21 families), and Eulipotyphla (599 species; 61 genera; 5 families), collectively making up 71.5% of total mammal species richness in MDD2. These orders are followed by Primates (522 species; 84 genera; 19 families), Artiodactyla (371 species; 138 genera; 23 families), and Carnivora (319 species; 129 genera; 16 families); all other mammal orders have fewer than 200 recognized species. Tubulidentata is currently the only mammal order with a single living species, since Microbiotheria now includes two species (although the exact species count has been hotly debated and ranges from one to three; D’Elía et al. 2016; Valladares-Gómez et al. 2017; Martin 2018; Suárez-Villota et al. 2018; Quintero-Galvis et al. 2022). The most speciose mammalian families are Muridae (876 species; 156 genera), Cricetidae (869 species; 162 genera), Vespertilionidae (535 species; 60 genera), Soricidae (488 species; 28 genera), Sciuridae (321 species; 64 genera), Phyllostomidae (231 species; 61 genera), and Pteropodidae (202 species; 46 genera). All other mammal families have fewer than 200 species, with 24 families having only 1 extant species (including Dugongidae and Thylacomyidae, which currently include 1 recently extinct and 1 extant species each). The three most speciose genera are *Crocidura* (220 species), *Myotis* (139 species), and *Rhinolophus* (114 species), which are also the only mammal genera exceeding 100 extant or recently extinct species; *Crocidura* exceeded 200 species and *Rhinolophus* exceeded 100 species as a result of new species recognized since MSW3.

Since MSW3, there have been 1,579 more species recognized as distinct (23.4% of all currently valid species), including 805 de novo species and 774 split species listed in MDD2, while 226 species have been lumped or removed, leading to a net increase of 1,353 species (+24.8%). Over this same 20-year period, at least one new species has been recognized in 20 out of 27 orders and 98 out of 167 families of mammals (this is likely to increase further as Pholidota/Manidae, which hasn’t had a new species recognized since MSW3, appears to have two additional species; Gu et al. 2023; Wangmo et al. 2025). The orders with the most species newly recognized since MSW3 are Rodentia (595 species; 21.7% relative to the order total in MDD2), Chiroptera (410 species; 27.6%), Eulipotyphla (166 species; 27.7%), and Primates (161 species; 30.8%). Of orders with over 100 species, Didelphimorphia (50 species; 39.7%) and Primates have the largest proportion of new species recognized since MSW3 relative to MDD2 totals; whereas Carnivora (50 species; 11.3%), Diprotodontia (18 species; 11.3%), and Artiodactyla (51 species; 13.8%) have the smallest proportion of new species recognitions. The families with the most new species recognized since MSW3 are Cricetidae (243 species; 27.7%), Vespertilionidae (173 species; 32.3%), Muridae (162 species; 19.9%), Soricidae (129 species; 26.4%), Phyllostomidae (76 species; 32.9%), and Didelphidae (50 species; 39.7%); all other families added fewer than 50 valid new species. Of families with over 100 total species, Didelphidae, Rhinolophidae (38 species; 33.3%) and Phyllostomidae have the largest proportion of new species recognized since MSW3.

Considering higher taxa, there has been a net increase of 123 recognized genera since MSW3 (+178 and –55), including 26 genera that appear to be de novo descriptions by having all included species described after MSW3 (*Alpiscaptulus*, *Antillomys*, *Baletemys*, *Calassomys*, *Chimaerodipus*, *Chingawaemys*, *Cordimus*, *Dryadonycteris*, *Drymoreomys*, *Eudiscoderma*, *Gracilimus*, *Halmaheramys*, *Hyorhinomys*, *Laonastes*, *Mictomicrotus*, *Mirzamys*, *Musseromys*, *Pattonimus*, *Paucidentomys*, *Pennatomys*, *Rungwecebus*, *Saxatilomys*, *Setirostris*, *Tonkinomys*, *Waiomys*, *Xeronycteris*). Diatomyidae is the only mammal family recognized since MSW3 where all extant species were described since MSW3 (originally named as a new family name, Laonastidae, but later shown to be a living representative of the Oligocene-Miocene fossil family Diatomyidae; Jenkins et al. 2005; Dawson et al. 2006).

### Regional Biodiversity

Across terrestrial biogeographic realms (Fig. 2A; Table 3), the Neotropical realm harbors the greatest number of currently recognized mammal species (1,839 terrestrial species), followed by the Afrotropical (1,505 species), Palearctic (1,143 species), Indomalayan (1,090 species), Australasian (860 species), Nearctic (716 species), and Oceanian (21 species) realms; the Antarctic biogeographic realm has no permanently terrestrially distributed mammal species. However, in terms of species density per land area, Indomalaya has the highest density of mammal species, followed by the realms of Australasia, Neotropic, Afrotropic, Nearctic, Palearctic, and Oceania (Table 3; note that the corrigendum to Burgin et al. 2018 corrected a previous error, also finding that Indomalaya has the highest density of mammal species; Burgin et al. 2019). As with species richness, the Neotropics also contain the most newly recognized species since MSW3 (563 species: 302 de novo, 261 split), followed by the realms of Afrotropic (294 species: 183 de novo, 111 split), Indomalaya (268 species: 144 de novo, 124 split), Palearctic (265 species: 92 de novo, 173 split), Australasia (131 species: 69 de novo, 62 split), Nearctic (114 species: 16 de novo, 98 split), and Oceania (3 species: 2 de novo, 1 split). The Neotropical biogeographic realm has the largest proportion of species that were recognized since MSW3 (30.6% of total species), while Oceania contains the smallest proportion (14.3%). Unlike other regional scales discussed below, MDD2 species totals for biogeographic realms do not incorporate marine-distributed species (all cetaceans, sirenians, pinnipeds, and marine mustelids), which encompasses 132 species (including 15 new species since MSW3), although 5 of these marine-distributed species also have inland populations that are included in biogeographic realm totals (specifically *Trichechus manatus*, *T. senegalensis*, *Pusa hispida*, *Orcaella brevirostris*, and *Neophocaena phocaenoides*).

**Figure 2.**
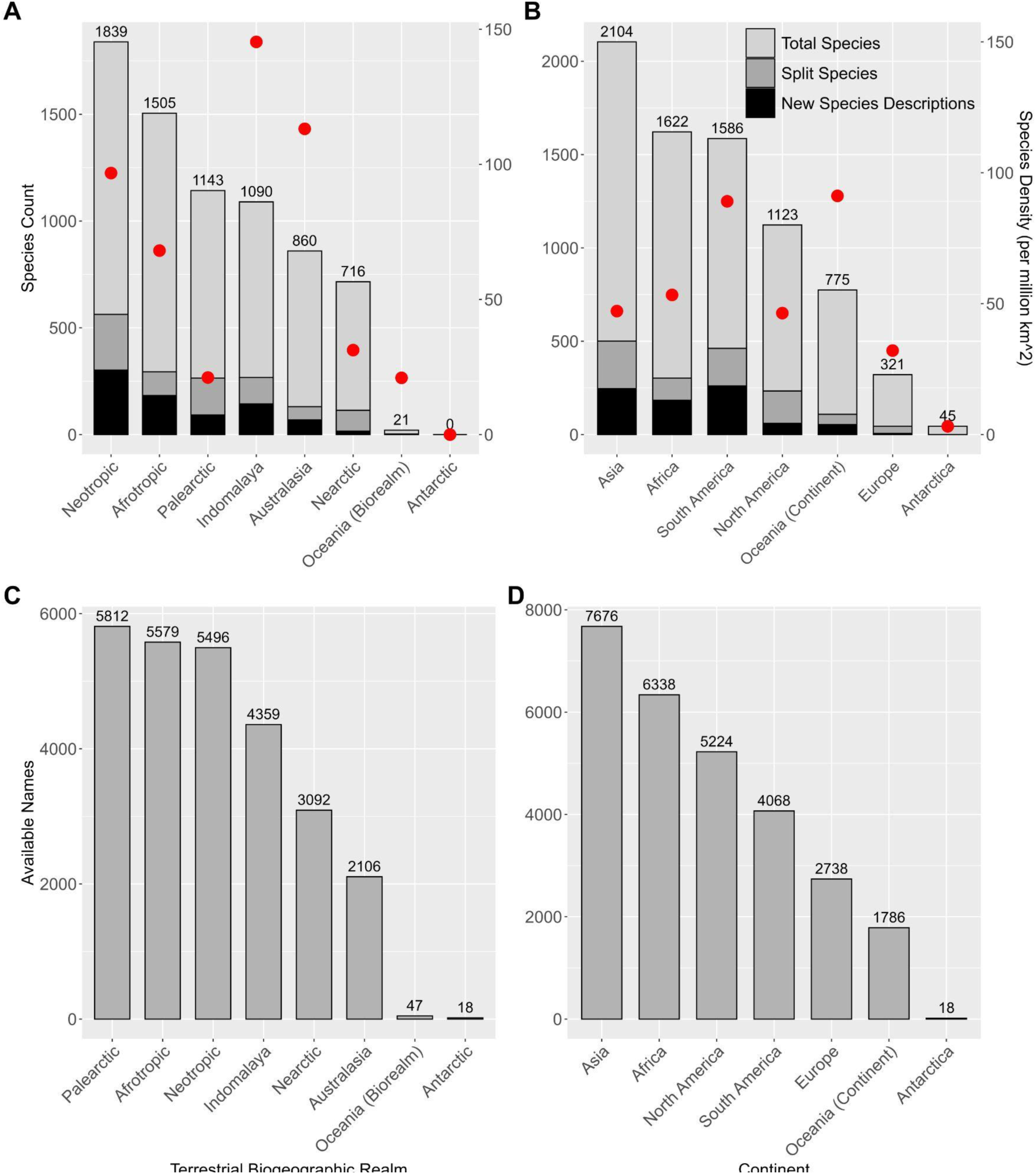
Summary of mammal species-rank taxonomy and nomenclature by terrestrial biogeographic realm and continent, including: A) total native, wild species (including recently extinct species) distributed within each region (excluding fully marine distributed species) with totals listed above each bar (broken into total species and species split or newly described since MSW3 was published), with total species density per region marked with red dots in species per million km^2^; B) total native, wild species distributed within and in the waters surrounding each continent with symbols given analogously to graph A; C) available names (including preoccupied, replacement, and suppressed names) with type localities within each terrestrial biogeographic realm with totals listed above each bar; and D) available names with type localities within each continent with totals listed above each bar. It should be noted that species are often distributed across multiple regions while available name type localities only appear in a single region. See Table 3 for numeric summaries of species richness and available name type localities relative to regional land areas.

**Table 3.** Summary of total species, available name type localities (including preoccupied, replacement, and suppressed names), and new species descriptions and split species since MSW3 found in each of the eight terrestrial biogeographic realms (not including marine species, other than for available name type localities) and seven continents (including marine species for all metrics), as well as the land area (in million km^2^) and species density (in species per million km^2^). Land areas for biogeographic realms come from Olson et al. (2001) (other than the Antarctic, which was recalculated to include all of continental Antarctica) and continents come from estimates listed on the Wikipedia pages of each continent. See Figure 2 for the graphical representation of this data.

| Region Name | Total Species<sup>a</sup> | Available Names | New Species Descriptions | Split Species | Land Area (million km<sup>2</sup>) | Species Density (per million km<sup>2</sup>)<sup>b</sup> |
| --- | --- | --- | --- | --- | --- | --- |
| **Biogeographic Realm** |  |  |  |  |  |  |
| *Neotropic* | 1,839 | 5,496 | 302 | 261 | 19.00 | 96.79 |
| *Afrotropic* | 1,505 | 5,579 | 183 | 111 | 22.10 | 68.10 |
| *Palaearctic* | 1,143 | 5,812 | 92 | 173 | 54.10 | 21.13 |
| *Indomalaya* | 1,090 | 4,359 | 144 | 124 | 7.50 | 145.33 |
| *Australasia* | 860 | 2,106 | 69 | 62 | 7.60 | 113.16 |
| *Nearctic* | 716 | 3,092 | 16 | 98 | 22.90 | 31.27 |
| *Oceania (Biorealm)* | 21 | 47 | 2 | 1 | 1.00 | 21.00 |
| *Antarctic* | 0 | 18 | 0 | 0 | 14.20 | 0.00 |
| **Continent** |  |  |  |  |  |  |
| *Asia* | 2,104 | 7,676 | 246 | 255 | 44.60 | 47.17 |
| *Africa* | 1,622 | 6,338 | 184 | 119 | 30.40 | 53.36 |
| *South America* | 1,586 | 4,068 | 261 | 201 | 17.80 | 89.10 |
| *North America* | 1,123 | 5,224 | 60 | 174 | 24.20 | 46.40 |
| *Oceania (Continent)* | 775 | 1,786 | 54 | 55 | 8.50 | 91.18 |
| *Europe* | 321 | 2,738 | 6 | 39 | 10.00 | 32.10 |
| *Antarctica* | 45 | 18 | 0 | 0 | 14.20 | 3.17 |

<sup>a</sup> Total Species includes recently extinct species but not domesticated or introduced species
<sup>b</sup> Species Density calculated as Total Species / Land Area

At the continental scale (Fig. 2B; Table 3), Asia contains the most recognized mammal species (2,104 species), followed by Africa (1,622 species), South America (1,586 species), North America (1,123 species), Oceania (775 species), Europe (321 species), and Antarctica (45 species). Of these, Oceania has the highest species density when weighted by continental land area, followed by South America, Africa, Asia, North America, Europe, and Antarctica (Table 3). Asia also has the highest number of newly recognized species since MSW3 (501 species; 246 de novo, 255 split), followed by South America (462 species; 261 de novo, 201 split), Africa (303 species; 184 de novo, 119 split), North America (234 species; 60 de novo, 174 split), Oceania (109 species; 54 de novo, 55 split), and Antarctica (0 species). The continent with the largest proportion of newly recognized species is South America (29.1% of total species), while the smallest other than Antarctica is Europe (14.0% of total species), closely followed by Oceania (14.1% of total species).

Between countries and offshore territories, the ten most speciose countries (Fig. 4A) are Indonesia (775 species), Brazil (768 species), China (671 species), Mexico (581 species), Peru (570 species), Colombia (520 species), Democratic Republic of the Congo (474 species), United States (468 species), Ecuador (438 species), and India (422 species). Many of these countries also occupy a similar placement in the amount of new species recognized since MSW3 (Fig. 4B), the top ten being Brazil (169 species; 22.0% of MDD2 species for that country), China (167 species; 24.9%), Peru (133 species; 23.3%), Indonesia (122 species; 15.7%), Colombia (120 species; 23.1%), Ecuador (118 species; 26.9%), Mexico (116 species; 20.0%), Vietnam (88 species; 25.6%), Madagascar (82 species; 32.3%), and Venezuela (72 species; 17.6%). Madagascar, Ecuador, and China have proportionately the most new species since MSW3 relative to total species in their country (see Fig. 4B). The number of endemic mammal species (i.e., found only in one country, including marine species), is highest in Indonesia (342 species; 44.1%), followed by Australia (286 species; 73.7%), Brazil (248 species; 32.1%), Madagascar (215 species; 84.7%), Mexico (209 species; 36.0%), China (156 species; 23.3%), Philippines (149 species; 63.4%), United States (123 species; 26.3%), Argentina (92 species; 23.2%), and Papua New Guinea (92 species; 31.2%). The countries with the highest proportion of mammal species endemism are Madagascar, Australia, Philippines, Indonesia, and Mexico (Fig. 4F).

**Figure 3.**
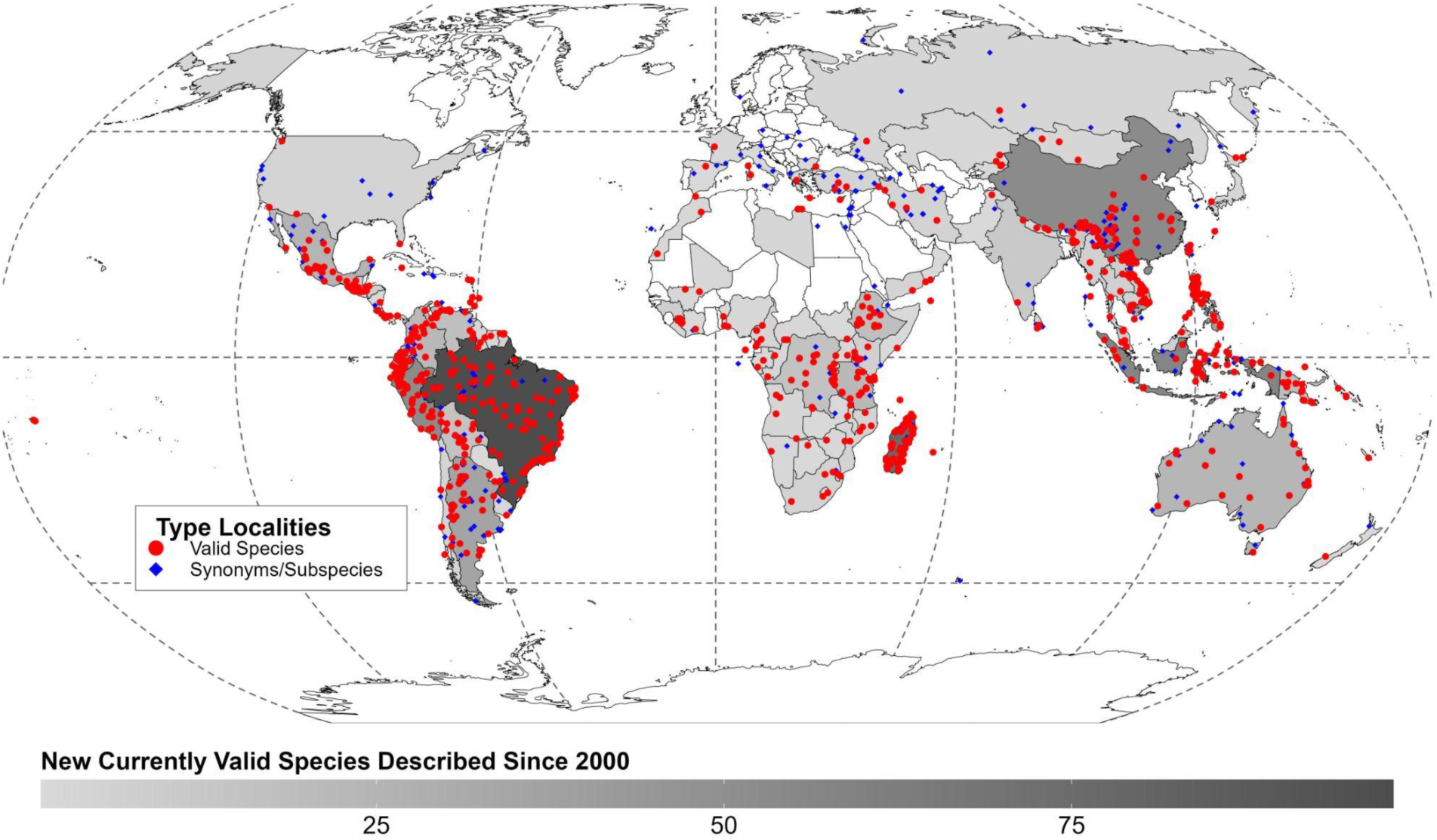
Type locality locations for species-rank mammal taxa described from 1 January 2000 to 15 August 2024. Map gray-scale country coloration represents the total number of currently valid species type localities found per country described within this period. Colored dots represent the exact georeferenced type locality of all currently valid species (red circles) and names currently considered synonyms and subspecies (blue diamonds) in MDD2. Most coordinates were generated from the original description of each name when included directly in publication, while others were georeferenced using GeoLocate or WikiMedia GeoHack place coordinates when not included in the original description publication. The georeferenced localities were mapped and both programmatically and visually vetted for accuracy. Latitude and longitude in decimal degrees is included for each of the names mapped here in the MDD2 synonym list, Supplementary Data SD2.

**Figure 4.**
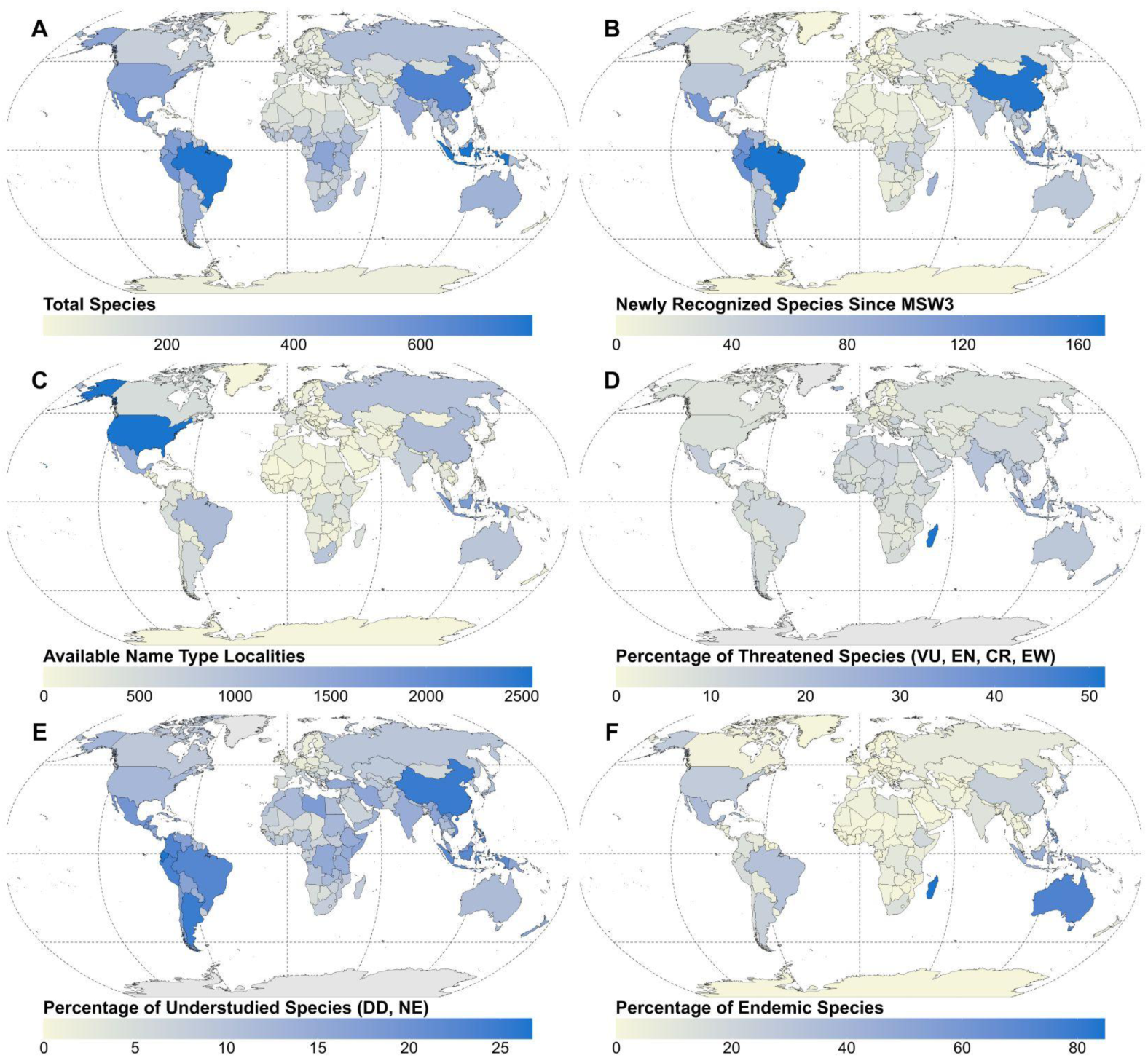
A) Total number of wild extant and recently extinct mammal species distributed within and in the waters surrounding each country and offshore territory. B) Total number of newly recognized species (both de novo and split) distributed within each country and offshore territory. C) Total number of available names (including preoccupied, replacement, and suppressed names) with type localities within each country and offshore territory. D) Percentage of species distributed in each country and offshore territory that are categorized as threatened (VU, EN, CR, EW) on the IUCN Red List (version 2024-1). E) Percentage of species distributed in each country and offshore territory that are either categorized as Data Deficient (DD) or Not Evaluated (NE) on the IUCN Red List, considered ‘understudied’ here. F) Percentage of species distributed in each country and offshore territory that are endemic to each region (only found in a single region).

Among states of the United States (Fig. 5), California has the most recognized mammal species (207 species), followed by New Mexico (150 species), Texas (148 species), Oregon (145 species), and Arizona (143 species). Hawai’i has the lowest species richness (28 species), having only 2 terrestrial and 1 non-cetacean marine mammal species recorded in the state (the rest being cetaceans); however, the terrestrial total will decrease by 1 in future MDD versions given recent evidence that all Hawaiian bat populations are attributable to *Lasiurus semotus* (Pinzari et al. 2023). The states with the most newly recognized species since MSW3 are California (20 species; 9.7% of total MDD2 species in state), Washington (18 species; 13.9%), Oregon (17 species; 11.7%), Texas (17 species; 11.5%), and New Mexico (17 species; 11.3%); by contrast, Alabama and the District of Columbia have only 1 species newly recognized since MSW3, and 11 other states only have 2 newly recognized species. The states with the most endemic species are California (17 species; 8.2%), Florida (5 species; 6.4%), Washington (4 species; 3.1%), and Alaska (4 species; 4.0%). See Supplementary Data SD7 for a numeric summary of nomenclatural data and species totals for all geographic regions, and Supplementary Data SD10 for a species matrix of country and offshore territory occurrences based on the MDD2 species list.

**Figure 5.**
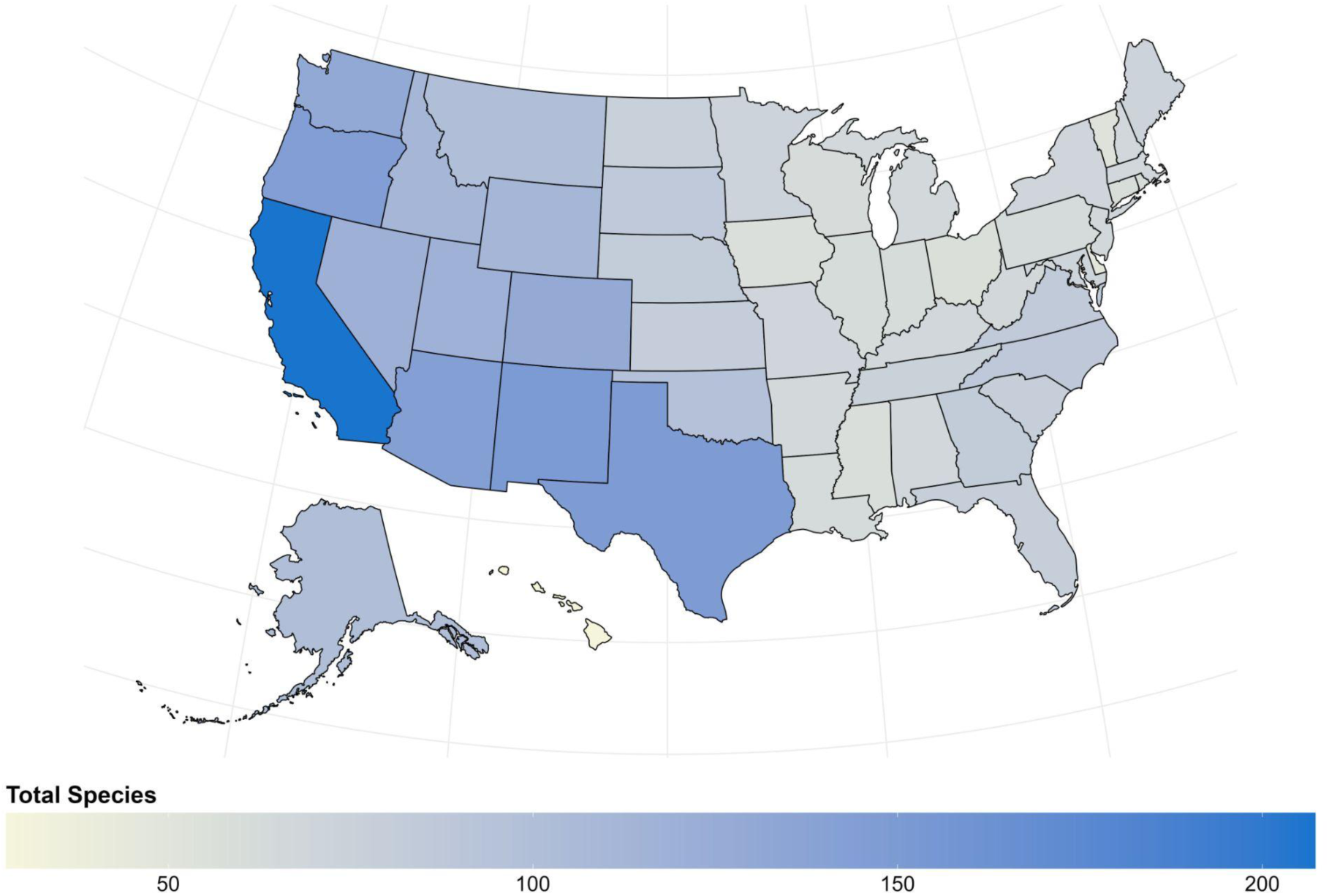
Mammal species richness across the 50 states of the United States, including both terrestrial mammals and marine mammals found along the coast of coastal states. State-level species totals are included in Supplementary Data SD7.

### Conservation Status

Of the 6,742 wild mammal species (excluding the 17 domestic species not assessed by the IUCN) recognized in MDD2, 14.0% (943 species) are Not Evaluated (NE) on the IUCN Red List because they are not recognized in the IUCN taxonomy (IUCN 2024a; see Fig. 6). The IUCN currently recognizes 5,983 mammal species, 5,799 of which are shared with MDD2 while 184 are considered by MDD2 to be synonyms of other species. Of the total species recognized in MDD2, 19.2% (1,295 species) are threatened with extinction (VU, EN, CR, and EW IUCN statuses) and 11.5% (773 species) are classified as Data Deficient (DD). Thus, fully 25.5% (1,716 species) of MDD2-accepted mammal species are either listed as DD or are NE relative to the IUCN Red List (collectively termed ‘understudied species’ later in this section, Fig. 4E, and Fig. 6). It should be noted that the total threatened species proportions listed above (and those later in this section) are calculated based on MDD2 totals, which include 943 wild NE species; however, the total proportion of threatened species would increase to 22.4% (1,339 species) and DD species to 14.0% (837 species) if considering only those MDD2 species listed on the IUCN and the IUCN species total of 5,983.

**Figure 6.**
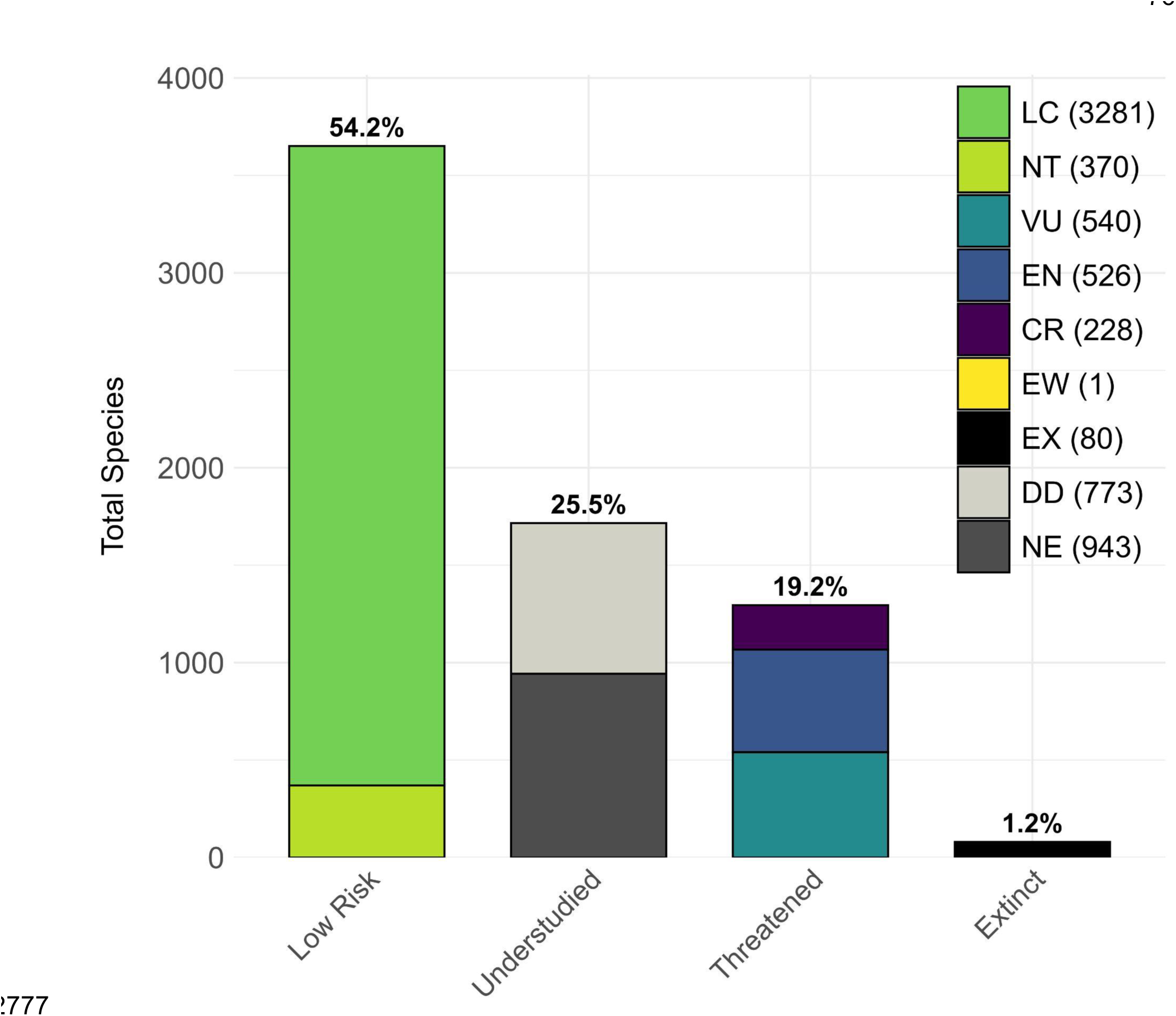
Proportion of the 6,742 wild mammal species (not including domestic species) recognized in MDD2 categorized by IUCN Red List category (version 2024-1): 54.2% (3651 species) are considered ‘Low Risk’ (Least Concern [LE] or Near Threatened [NT]); 25.5% (1716 species) are considered ‘Understudied’ (Data Deficient [DD] and Not Evaluated [NE]); 19.2% (1295 species) are considered ‘Threatened’ (Vulnerable [VU], Endangered [EN], Critically Endangered [CR], or Extinct in the Wild [EW—not visible]); and 1.2% (80 species; note that some MDD2 extinct species are NE on the IUCN) are considered globally Extinct (EX) on the IUCN. Total species per category are listed next to the category abbreviation in the figure legend while percent of total MDD2 species per column appears above each column.

MDD2 considers 113 species to be recently extinct and documents the extinction of 25 entire genera and 7 entire families since the year 1500. In contrast, the IUCN lists 84 recently extinct species, 79 of which are shared with MDD2, leaving 32 recently extinct species listed in MDD2 that are NE by the IUCN, and 5 recently extinct species in the IUCN not recognized by MDD2 due to taxonomic revisions (e.g., *Cryptonanus ignitus* is now a synonym of *C. chacoensis*; Teta and Díaz-Nieto 2019). Furthermore, 2 species considered extinct in MDD2 are currently considered extant by the IUCN (*Dasycercus cristicauda* as NT; *Tonatia saurophila* as LC) despite taxonomic revisions elevating the extant populations to valid species under different names and restricting the original name to extinct populations (Basantes et al. 2020; Newman-Martin et al. 2023).

The orders with the largest number of threatened extant species are Primates (335 species; 64.2% of total MDD2 species), Rodentia (313 species; 11.4%), Chiroptera (217 species; 14.6%), and Artiodactyla (122 species; 32.9%). Of orders with over 100 total species, Primates have the highest proportion of threatened species, followed by Artiodactyla, Diprotodontia (51 species; 32.1%), and Carnivora (75 species; 23.5%). All living species of Pholidota (8 species), Proboscidea (3 species), and Sirenia (4 species, plus 1 extinct) are considered threatened by the IUCN. The families with the most threatened species are Muridae (120 species; 13.8%), Cercopithecidae (114 species; 69.1%), Cricetidae (83 species; 9.5%), and Pteropodidae (73 species; 35.6%). Of families with over 100 species, Cercopithecidae, Pteropodidae, Bovidae (50 species; 32.1%), and Rhinolophidae (18 species; 15.8%) have the largest proportion of threatened species. There are 18 families where every species in the family is considered threatened by the IUCN, the most speciose of which are Lepilemuridae (25 species), Lemuridae (21 species), Hylobatidae (20 species), and Indriidae (19 species); note that all four of these families are within Primates, three being Madagascar endemic lemurs. Another 18 families have 50% or more of their species considered threatened on the IUCN.

Considering the most understudied species per clade relative to IUCN status, the orders with the highest proportion of DD species are Scandentia (5 species; 21.7% of total MDD2 species), Macroscelidea (4 species; 20%), and Cingulata (5 species; 20%), while Rodentia (371 species; 13.5%), Chiroptera (215 species; 14.4%), and Eulipotyphla (105 species; 17.5%) have the highest raw numbers of DD species. The orders with the highest proportion of MDD2-accepted species that are NE by the IUCN are Microbiotheria (1 species; 50%), Pilosa (7 species; 41.2%), and Didelphimorphia (39 species; 30.9%); whereas, Rodentia (458 species; 16.7%), Chiroptera (184 species; 12.4%), and Eulipotyphla (113 species; 18.9%) have the highest raw numbers of NE species. Combining these DD and NE categories, the orders with the most understudied species are Rodentia (829 species; 30.2%), Chiroptera (399 species; 26.9%), and Eulipotyphla (218 species; 36.4%). Only 8 of 27 orders and 72 of 167 families of mammals are currently ‘completely evaluated’ by the IUCN, meaning that all MDD2-accepted species in that family are ranked in categories other than DD or NE.

Across terrestrial biogeographic realms, there are 378 NE species distributed in the Neotropical (20.6% of total MDD2 species in that region), 196 in the Palearctic (17.2%), 135 in the Indomalayan (12.4%), 122 in the Afrotropical (8.1%), 94 in the Nearctic (13.1%), 63 in the Australasian (7.3%), and 1 in the Oceanian (4.8%). Similarly, the number of total understudied species is highest in the Neotropical (627 species; 34.1%), followed by the Afrotropical (306 species; 20.3%), Palearctic (293 species; 25.6%), Indomalayan (287 species; 26.3%), Australasian (159 species; 18.5%), Nearctic (107 species; 14.9%), and Oceanian (2 species; 9.5%). The most species threatened with extinction are located in the Afrotropical (348 species; 23.1%), followed by the Indomalayan (283 species; 26.0%), Neotropical (237 species; 12.9%), Australasian (214 species; 24.9%), Palearctic (127 species; 11.1%), Nearctic (92 species; 12.9%), and Oceanian (10 species; 47.6%). The biogeographic realms with the most recently extinct species in MDD2 are the Neotropical (46 species, including 37 in the Caribbean) and the Australasian (39 species, including 33 species in Australia), with all other regions including 8 or less recently extinct species.

Among continents, South America has the most species that are NE by the IUCN (290 species; 18.2%), followed by Asia (284 species; 13.5%), North America (192 species; 17.1%), Africa (130 species; 8.0%), Oceania (55 species; 7.1%), Europe (32 species; 10.0%), and Antarctica (0 species). However, Asia has the overall most understudied species (549 species, 26.1%), followed by South America (519 species, 32.7%), Africa (328 species, 20.2%), North America (241 species, 21.5%), Oceania (122 species, 15.7%), Europe (38 species, 11.8%), and Antarctica (2 species, 11.8%). The continent with the most species threatened with extinction is Asia (432 species; 20.5%), followed by Africa (363 species; 22.4%), South America (205 species; 12.9%), Oceania (178 species; 23.0%), North America (147 species; 13.1%), Europe (42 species; 13.1%), and Antarctica (6 species; 13.1%). The continents with the most recently extinct species in MDD2 are North America (41 species, including the same 37 species in the Caribbean as in the Neotropical biogeographic realm) and Oceania (40 species, including the same 33 species in Australia as in the Australasian biogeographic realm), with all other continents including 11 or fewer recently extinct species. These figures highlight the high totals of recent mammal extinctions in the Caribbean and Australia (see Woinarski et al. 2015; Cooke et al. 2017).

Among countries and offshore territories, the most understudied species (Fig. 4E) are in Brazil (174 species; 22.7% of MDD2 total for that country), Indonesia (171 species; 22.1%), China (168 species; 25.0%), Peru (136 species; 23.9%), Colombia (121 species; 23.3%), Ecuador (117 species; 26.7%), Mexico (113 species; 19.5%), Argentina (99 species; 24.9%), Vietnam (73 species; 21.2%), and Democratic Republic of the Congo (72 species; 15.2%). Ecuador, China, and Argentina contain the highest proportion of understudied species relative to the MDD2 total for that country. The greatest concentrations of species threatened with extinction (Fig. 4D) are found in Indonesia (211 species; 27.2%), Madagascar (131 species; 51.6%), Brazil (94 species; 12.2%), Mexico (88 species; 15.2%), India (82 species; 19.4%), China (77 species; 11.5%), Malaysia (77 species; 22.0%), Australia (68 species; 17.5%), Thailand (63 species; 18.9%), and Colombia (57 species; 11.0%). The countries with the highest proportion of threatened species relative to the MDD2 total are Madagascar, Solomon Islands (24 species; 30.4%), and Mauritius (7 species; 29.2%). The countries and offshore territories with the most recently extinct species recognized in MDD2 are Australia (33 species), Dominican Republic (14 species), Haiti (14 species), Cuba (8 species), and Madagascar (6 species), further highlighting the recent extinctions in Australia and the Caribbean. A total of 44 countries and offshore territories have had at least one species go extinct that was partly or wholly distributed within its borders since 1500 CE.

### Nomenclatural Metadata

MDD2 contains a curated dataset of 50,230 species-rank names that are applicable to all currently recognized mammal species (including *nomina dubia* and *species inquirendae* that were meant to apply to extant or recently extinct mammals when described). In terms of nomenclatural status, these names consist of 28,382 available names (including 925 available *nomina nova*, 1,100 preoccupied names, and 39 suppressed names; this total includes all names initially described as available and those that were subsequently made unavailable due to being preoccupied or suppressed), 2,127 unavailable names (including all names used with the intent of being available for taxonomic purposes that are unavailable from publication according to the ICZN Code), 3,445 spelling variants, 16,182 name combinations, and 435 instances of other subsequent name use (Table 4; see also Supplementary Data SD8). There are 23,973 more species-rank names listed in MDD2/Hesperomys compared to MSW3, which listed 26,257 such names. This difference is primarily due to the inclusion of more unavailable names, spelling variants, and name combinations (i.e., names not usable for taxonomic purposes), although a number of available names described subsequent to, or omitted from, MSW3 have been added to the MDD2/Hesperomys synonym list as well.

**Table 4.**
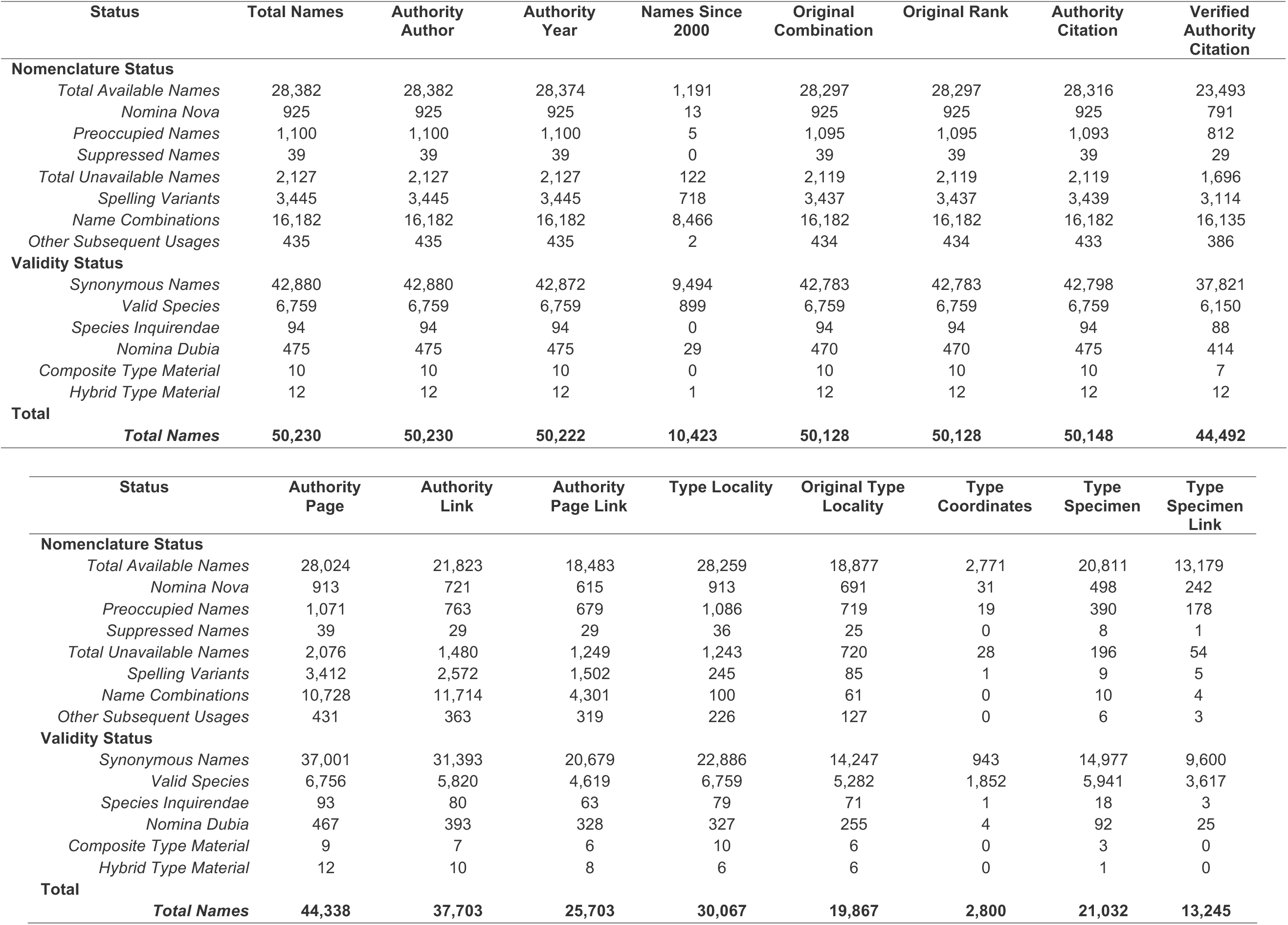
Summary of nomenclatural metadata released in MDD2 separated into nomenclatural status and validity status. Totals for available names include preoccupied names, *nomina nova*, and suppressed names (all of which are also summarized separately). Totals for unavailable names include all nomenclatural status conditions that are not some form of subsequent use, while name combinations, spelling variants, and other miscellaneous subsequent uses are summarized separately (see Supplementary Data SD8 for a full list of nomenclatural and validity statuses and finer scale numeric summaries). Multiple nomenclatural labels can be applied to a single name, while only one validity status can be applied to a name, so column totals summarize the validity statuses.

Authority authors are included for all names and valid species in MDD2, but authority year is missing for 8 names (all are synonyms of *Equus* species, 7 of which have been found after the MDD2 cutoff date and will be included in later versions). For nearly all synonymous names and every valid species, MDD2 includes original name combination, taxonomic rank, and some degree of citation information (99.8% of all names). Citation information has been fully verified for 88.6% of names (91.0% of valid species), and the page number on which the name first appears or is described is included for 88.3% of names (and all but 3 valid species). For 75.1% of names (86.1% of valid species) a hyperlink to the full text of the original description was located, and for 51.2% of names (68.3% of valid species) direct links were included to the page that name was first mentioned or described on the BHL.

Type localities were also recorded for almost all applicable names (99.6% of originally available names and all valid species) and have been verified in the original description for 66.5% of available names and 78.2% of valid species. MDD2 also includes coordinates of the type localities in decimal degrees for 9.8% of available names and 27.4% of valid species (including almost all names described after the year 2000). Type specimen locations were gathered for 73.3% of originally available names (87.9% of valid species), 46.4% of which (53.5% of valid species) are linked to the specimen’s database entry in an online collections catalog. Most species descriptions designate a single type specimen (the holotype), which MDD2 records for 18,396 names. However, there are also 1,111 names based on multiple type specimens (syntypes) and 1,092 for which a lectotype has been selected from among a series of syntypes. There are 144 cases where the original type material has been lost or was nonexistent in the first place, resulting in a neotype being designated. There are also 289 names in MDD2 recorded as having nonexistent type specimens, which will require future neotype designation. We estimate that MDD2’s coverage is nearly complete for type specimens housed in North America, where almost all major institutions have published type catalogs. Most of the names for which we have not yet recorded type specimen information likely have no surviving types or were based on types housed in a European collection.

### Historical Taxonomy and Nomenclature

Species-rank available names were described from 1758 to 2023 at an average rate of 106.2 names per year (Fig. 1; raw data and running means available in Supplementary Data SD9). However, when divided into roughly 50-year time bins, the following average rates of yearly naming emerge: 32.7 (1758–1799), 95.0 (1800–1849), 117.8 (1850–1899), 248.4 (1900–1949), 53.1 (1950–1999), and 48.5 (2000–2023). The time bins with the lowest naming rates were 1758–1799 and 2000–2023, although this latter time bin is about half as long, while the highest-rate periods were 1900–1949 and 1850–1899. From 1758–1800, there were 19 years where fewer than 10 available names were described and four years where none were described (1759, 1762, 1763, 1764; although 1762 has unavailable names). Sustained rates of species-rank name description were not reached until 1819, after which every year had >30 available name descriptions until 1973. Yearly peaks associated with single publications occurred in 1827 (206 names), 1842 (287 names), 1860 (255 names), and 1867 (238 names). During the 1890s, most years had >200 described names, peaking at 366 in 1898. This trend continued into the first decades of the 20th century, leading to 1913 as the year with the most documented new name descriptions across the ∼267 years of taxonomic history (529 available names described). The mid-20th century experienced a sustained decrease in naming rates, with only three years since 1943 having more than 100 available name descriptions (1946 with 120, 1955 with 127, and 1956 with 108). Only 24 names were described in 1973, making it the year with the least name descriptions since the early 19th century (only 12 of which are currently valid species). The proportion of names that are currently recognized as valid species in MDD2 increased from the 1970s to the 21st century as rates of subspecies name description decreased. The year 2021 has the most names described during the 21st century (69 names) and 2001 had the least (33 names). There have been 28 species-rank names described in 2024 prior to the MDD2 cutoff date of August 15th (including 24 currently valid species; increasing to 42 valid species during the writing of this manuscript).

Considering the history of taxonomic rank use over time, most species-rank names described before the 20th century were described as species rather than subspecies, forms, or varieties (Fig. 1; forms and varieties before 1900 were often equivalent to the subspecies rank and are treated equivalently here). Over 70% of names were described initially as species until the time bin of 1895–1904, after which it steadily fell until a resurgence in the 21st century. While species were the primary taxonomic unit used for name descriptions during the 18th and 19th centuries, there were significant jumps in subspecific name descriptions (generally as forms or varieties) in 1788 (61.4% of available names described that year were subspecies), 1792 (62.0%), 1801 (73.9%), and 1860 (69.8%). The early 20th century marked an increase in the prevalence of polytypic species (species with multiple subspecies), with >50% of all names described as subspecies from 1922 to 1973, peaking in 1948 at 89.0%. Following this period, the prevalence of subspecies name descriptions began to decrease, dropping below 50% in 1974. This time period lasted until 1999 when subspecies descriptions consistently dropped to below 50% of all yearly name descriptions. Subspecies descriptions became considerably less common after 2003, decreasing to only 6.4% of names described in 2014 and not again exceeding 25% of yearly names.

The majority of all available mammal names are considered synonyms of valid species on MDD2 (i.e., 21,254 names; 74.9% of available names). Of those, 17,906 names (63.1%) were initially described at the species rank while 10,367 (36.5%) were initially described as subspecies, forms, or varieties. Furthermore, 11,508 names (40.6%) described as species have been lumped, while 698 names (2.5%) described as subspecies have been subsequently split.

There are 6,055 names (21.3%) described as species that are recognized as valid species in MDD2, although it should be noted that this total does not consider that many of these names have been treated as synonyms or subspecies before being revalidated later.

Most names that were described as species and since lumped (i.e., now synonymous names in MDD2) were published before 1920, with the 10-year running mean of this statistic consistently over 50% from 1785 to 1920. From 1927 to 2023 (other than 1976 and 2001), fewer than 25% of the available species-rank names described per year were lumped. In contrast, the rate of taxa that were initially described as subspecies that have since been elevated to species (i.e., split out from another valid species and now recognized in MDD2) was highest throughout the 20th century, with a 10-year running mean of ∼1–4% from 1886 to 1958 compared to ∼5–9% from 1959 to 2001. Few years before 1886 and after 2009 include names described as subspecies that are now recognized as species, with most yearly rates ranging from ∼0–1% during these time periods. It should be noted that these rates are based on the taxonomic rank at which each name was originally described and whether it is now considered a synonym or valid species in MDD2. Thus, these values do not capture when each name was split or lumped or whether a name was split or lumped multiple times throughout their taxonomic history.

Across terrestrial biogeographic realms, the Palearctic has the most available names with type localities in that region (20.5% of total available names), followed by the Afrotropical (19.7%), Neotropical (19.4%), Indomalayan (15.4%), Nearctic (10.9%), Australasian (7.4%), Oceanian (0.2%), and Antarctic (<0.1%) (Fig. 2C; Table 3). The continent with the most available names is Asia (27.1%), followed by Africa (22.3%), North America (18.4%), South America (14.3%), Europe (9.7%), Oceania (6.3%), and Antarctica (<0.1%) (Fig. 2D; Table 3). The ten countries with the most available name type localities are the United States (2,552 names), Indonesia (1,732), Mexico (1,301), Brazil (1,088), China (1,084), Russia (975), South Africa (916), Australia (895), India (689), and Kenya (523) (Fig. 4C).

Type localities of new species-rank names described since 2000 have a global distribution, but there is a strong latitudinal gradient, with a larger proportion of names described from tropical over temperate regions (Fig. 3). Species-rank name type localities are concentrated throughout high-elevation regions (e.g., Andes, East African and Ethiopian highlands, and Himalayas), islands (e.g., Madagascar, the Philippines, and Sulawesi), and in biodiversity hotspots (e.g., Atlantic Forest). Conversely, there appear to be gaps or lower concentrations of new type localities from regions that may be expected to have similarly unexplored mammal faunas, such as Central and South Asia, West and Central Africa, portions of the Amazon Basin, and various Indonesian islands (e.g., Java, Sumatra, Borneo). Few newly described names from the 21st century have type localities in northern North America, Europe, and northern Asia, and most that do are either subspecific names or names now regarded as synonyms. The 5 countries with the most type localities of valid species described during the 21st century are Brazil (98 names), Madagascar (78), Indonesia (57), China (53), and Peru (52).

### Website and Database Usage

The MDD website located at https://mammaldiversity.org increasingly serves a global audience of users. Over a recent 17-month period, it was visited by 1,200–2,900 users/month for a total of 25,991 unique users that were located in 170 different countries (Google Analytics: 19 May 2023 to 19 October 2024). The countries with the highest overall usage during this period were the United States (34.3% of users), Brazil (6.7%), Mexico (6.2%), and China (5.4%), which topped the 12 countries with over 500 users and the 50 countries with over 50 users (see Supplementary Data SD11 for complete usage data). Global usage rates rose from an average of ∼450 weekly users (months 1–10 of tracking) to ∼700 (months 11–17). A large spike in MDD website traffic on 15-17 July 2024 involving 1,326 weekly users likely stemmed from the release of the MDD v1.13 taxonomy on 13 July 2024. The archived versions of the MDD taxonomy that are hosted on Zenodo at https://doi.org/10.5281/zenodo.4139722 have garnered 33,208 total page views and 17,159 downloads over the past four years (Sep 2020–Oct 2024).

## DISCUSSION

The Mammal Diversity Database version 2.0 (MDD2) is a digital compendium of global mammal taxonomic, nomenclatural, and geographic information that is more complete and accessible than prior efforts, standing as a stepping stone on the path to greater knowledge about mammals. MDD2 provides an online overview of taxonomic and distributional studies authored by mammalogists worldwide over the past 267 years (1758–2024), jointly illuminating the latest advances in taxonomic understanding of this charismatic vertebrate clade. The release of MDD1 seven years ago marked a shift from decadal printed volumes to semi-annual online versions (Burgin et al. 2018). However, continued advances to the MDD have required improved strategies to find, extract, and link relevant information from the disparate sources needed to assemble real-time updates to global mammal taxonomy. Because much of the information used to update the MDD is in rare archives, offline sources, or behind paywalls, manual data curation by a dedicated team of students and volunteers has been crucial in collating these data and linking their contained biodiversity knowledge.

### Documenting 267 Years of Change: Historical Perspectives on Mammalian Taxonomy

The curatorial efforts of MDD2 and Hesperomys have uncovered extensive information relevant to the taxonomic and nomenclatural history of mammals, such as type localities, type specimens, and original description citations and textual links. These data remain important for modern taxonomic revisions when considering the identity and status of previously published names but also paint a broad trajectory of mammalogical study spanning from the dawn of Linnaean taxonomy to the present. The MDD2 dataset allows for novel explorations of how the recognized taxonomy of mammal species and higher taxa has changed through time, which is briefly summarized here.

The history of mammalian taxonomic and nomenclatural descriptions shows an arc of changing biodiversity knowledge from 1758 to 2024 (Fig. 1). This history is here broken into four major eras: (i) 1758–1880, when many large, widespread, and charismatic mammals were described sporadically in monographs; (ii) 1881–1939, the peak of descriptive taxonomy associated with greater use of subspecies and journal-based, taxon- or region-specific publications; (iii) 1940– 1999, a period of decreasing taxonomic descriptions, peak use of subspecies, and transition toward revisionary taxonomy; and (iv) 2000-present, the current period of technology-driven research advances in integrative revisionary mammalian taxonomy. We discuss each of these historical eras in the following sections, leading to the current era, which is discussed in a separate section (see *Mammalian Systematics in the Modern Era*).

*(i) 1758–1880*. The late 18th century featured the first publications to describe binomial names for many charismatic mammal species (e.g., ungulates, carnivores, cetaceans; particularly in Europe), setting the foundation for the development of modern taxonomic and nomenclatural practices. Binomial nomenclature and taxonomy for mammals and other animals starts with C. Linnaeus in Sweden (1758), whose *Systema Naturae* (10th edition) still holds the title of proposing the largest number of currently valid mammal species in a single publication. These species were initially placed in 8 orders, but only Primates is still in use today, and 39 genera, all of which are still in use except *Simia*, which is suppressed for nomenclatural use (ICZN 1929). During this period, there was little standardization in the use of taxonomic units other than orders, genera, and species, with many genera having species compositions equivalent to modern mammal families, suborders, or orders (e.g., *Vespertilio* was used for all bats by Linnaeus). The use of families as a taxonomic grouping between orders and genera would not be applied to mammals until G. Cuvier’s (1797) *Tableau Élémentaire de l’Histoire Naturelle des Animaux* and would not be given the consistent suffix ‘-idae’ in mammals until J.E. Gray (1821). There are considerably fewer species-rank names described from this period compared to the 19th and 20th centuries, although there are two notable single-year spikes in species descriptions associated with single publications: Linnaeus (1758) with 208 names (152 currently valid species) and R. Kerr in Britain (1792) with 202 names (27 currently valid species). Other foundational authors and publications describing mammal species richness during the late 18th century include P.S. Pallas in Russia (1766, 1767–1774), and J.C.P. Erxleben (1777), J.F. Gmelin (1788), and J.C.D. von Schreber in Germany (1774–1844; some sections published posthumously). Most of these publications were monographs adding to or revising previously published global animal or mammal taxonomies rather than journal articles describing individual species or clades.

During this period, many species were renamed or described multiple times across publications, resulting in many synonymous names for single species. Some names were even described from the same type material or just to replace previously published names with the authors’ own (e.g., *Mandrillus sphinx* having six available names in the 1700s alone). Species-rank names described during this period were often based on illustrations or firsthand accounts of living or recently deceased animals without reference to voucher specimens, but museum specimens would become the preferred reference material for biological descriptions later in the 19th and early 20th century (i.e., museum-based type material). Early in the 19th century, taxonomic works were mostly published in Europe, involving prolific authors such as É. and I. Geoffroy Saint-Hilaire, G. and F. Cuvier, and A.G. Desmarest in France, J.E. Gray and G.R. Waterhouse in Britain, L. Fitzinger in Austria, and J.A. Wagner and W.C.H. Peters in Germany. However, through the 19th century, some researchers of European heritage working elsewhere started publishing taxonomic literature, including S.F. Baird, C.S. Rafinesque, and E. Coues in the United States, A. Smith in South Africa, and E. Blyth and B.H. Hodgson in India. Although publications through the 19th century were moving away from large monographs and toward journal articles, some monographic works still contributed to major individual peaks in taxonomic activity during four major single-year peaks in species-rank name descriptions from 1800–1890: those of R.P. Lesson in France (1827, 1842) and Fitzinger (1860, 1867; although most of Fitzinger’s names were domestic breeds proposed as forms or varieties).

The 19th century also saw significant philosophical advances in evolutionary theory, which drove further advances in the understanding of mammal biodiversity. The higher classification of mammals began to take shape with H.M.D. de Blainville, who used reproductive biology and skull and skeletal features to group mammals into distinct subclasses: monotremes (Ornithodelphes), marsupials (Didelphes), and placentals (Monodelphes) (de Blainville 1816, 1864). These works paralleled the evolutionary ideologies expressed in C. Darwin’s *Origin of Species* (1859) through the emphasis of skeletal similarities and adaptive alterations among related taxa, which were further emphasized by other contemporary works focused on the classification of mammals and other animals based on tooth morphology (Giebel 1855), brain structure (Bonaparte 1840), and reproductive biology (Owen 1868). The modern terms for these three extant mammal clades were established by T.N. Gill (1872), who initially proposed Prototheria for monotremes and Eutheria for marsupials and placentals, and T.H. Huxley (1881), who then termed Metatheria for the marsupials (albeit not initially intended as a scientific name), leaving placental mammals in Eutheria. The terms Prototheria, Metatheria, and Eutheria have varied in definition both historically and in modern use, but are usually given a broader cladistic definition, respectively including the living and related stem group fossil relatives (e.g., de Queiroz and Gauthier 1992) of the (i) monotremes (now often included under the clade Australosphenida, treating Prototheria as a defunct paraphyletic clade name), (ii) marsupials (often called Marsupialia for the crown clade), and (iii) placentals (often called Placentalia for the crown clade). Prototheria, Metatheria, and Eutheria are retained in the MDD to match the most common use of the names of extant mammals for the time being. A more detailed account of the early history of mammalian taxonomic research is available in Burke (1968).

*(ii) 1881–1939*. Prior to the 1890s, taxonomic and nomenclatural practices were unstandardized, leading to differential naming systems and confusion regarding a given species’ proper name, especially as many authors knowingly renamed already-named species. The lack of structure spurred the need for standardized nomenclatural and taxonomic rules for zoological taxonomy, which culminated in the publication of the *International Rules of Zoological Nomenclature* in 1905 (Blanchard et al. 1905) that would eventually develop into the first edition of the ICZN Code in 1961 (ICZN 1961). These documents established fundamental nomenclatural rules that are still used today, including the principle of priority for scientific names, the use of physical type material and type localities to define species-rank names, and the rule that names must be published according to the ICZN Code to be considered available. Thus, this period is further characterized by the appearance of more seriously cataloged museum type specimens and more precisely defined type localities (rather than broadly defined type localities at the country or continent level).

This standardization contributed to the rapid increase in mammalian taxonomic activity during the late 19th and early 20th century, which was also characterized by an increasing prevalence in the use of the subspecies rank (replacing forms and varieties). Mammal-focused expeditions also occurred more frequently, generally associated with colonial exploration, resulting in specimens being sent to museums in Europe and North America (rarely in the country of collection; e.g., see Mohammed et al. 2022). These trends continued into the first decades of the 20th century, expanding to researchers of European heritage in Germany (e.g., P. Matschie), the United States (e.g., C.H. Merriam, J.A. Allen, W.H. Osgood, G.S. Miller), Russia (e.g., S.I. Ognev), China (e.g., P.M. Heude), and South Africa (e.g., A. Roberts). Additionally, some of the first researchers of non-European heritage started publishing mammal names, such as Yang Zhongjian (then known as C.C. Young) from China and K. Kishida and T. Mori from Japan (Japanese was also the first non-European language known to feature a mammal name description). However, this rapid increase is most prominently associated with O. Thomas and other researchers at the British Museum of Natural History. Thomas, who initially wanted to study echinoderms, was the most prolific mammalian taxonomist in history, describing ∼2,900 genus- and species-rank names from 1880–1929 (Hill 1990), including 2,725 available species-rank names (9.6% of available names), 937 of which are currently valid species in MDD2.

By the 1920s, new taxonomic descriptions usually appeared in articles focused on individual taxa, regions, or expeditions published in journals instead of monographic books. A single journal, the *Annals and Magazine of Natural History*, published the descriptions of 3,434 available species-rank mammal names from 1841–1959, 12.1% of available names in MDD2 (including 990 currently valid species). Other prolific journals during and after the late 19th century include the *Proceedings of the Biological Society of Washington* with 1,500 available species-rank names from 1886–2021 (more consistently before the 1970s) and the *Proceedings of the Zoological Society of London* with 1,483 available species-rank names from 1831–1962. Additionally, the American Society of Mammalogists, and the associated *Journal of Mammalogy* (including 616 available species-rank names), was founded in 1919, establishing a nexus for focused mammalogical research in the United States and eventually the Americas more broadly.

A large proportion of the names proposed as species during this time period are now treated as subspecies or synonyms (Fig. 1, upper panel). A prime example of the frequency with which now-synonymous names were proposed during this period is Merriam’s (1918) treatment of the Brown Bear (*Ursus arctos*) as ∼74 distinct species in North America alone, as opposed to the currently defined single Holarctic species. Researchers primarily described mammal species based on relatively few specimens sent to museums from widely separated localities.

Additionally, species were often described based on subtle morphological distinctions, and some of the traits used for species diagnoses at this time are now known to vary substantially within species relative to environmental factors (e.g., pelage coloration). These dynamics contributed to the description of many names initially proposed as species or subspecies that are now recognized as a single species with intraspecific variation (e.g., over 200 available synonymous names referable to *Thomomys bottae*, as currently defined). Furthermore, most of the names described during this period were not actively cataloged in global-scale compendia (other than E.-L. Trouessart’s series cataloging living and fossil mammals earlier in the period; Trouessart 1897, 1899, 1905). In retrospect, this practice of not cataloging all described names has made it difficult to estimate how many mammal species were actively recognized across different time periods, particularly so in the early 20th century. Thus, subsequent efforts were needed to both catalogue and review the influx of new names, ushering in the next era in mammalian taxonomy and biodiversity research, which was characterized by a shift from descriptive to revisionary taxonomy.

*(iii) 1940–1999*. Descriptive taxonomic activity began decreasing after the 1920s, and especially into the late 1930s and 1940s following the Great Depression and leading into World War II. During this period, fewer names were being described, and even fewer are now considered valid species (e.g., only 3 currently valid species were described in 1961). In contrast to the previous period, proportionately more new names described during the mid-to late 20th century were described as subspecies, and many names described earlier as species were joined together into polytypic species with many subspecies. This decrease in species-rank name descriptions likely stems from many factors, including global political instability that shifted research funding away from systematics. There were also significant advances in the philosophy of species concepts, evolutionary theory, availability of specimens, and molecular methods during this period that contributed to a shift from descriptive to revisionary taxonomy.

To organize the large number of names coined in previous decades, several major regional compendia were published, such as Allen (1939) for Africa, Ellerman and Morrison-Scott (1951) for Eurasia, Miller and Kellogg (1955) for North America, and Cabrera (1958, 1961) for South America, among others. Although the taxonomic information in these treatises is now dated, they still remain invaluable as sources of bibliographic and geographic information for mammal names historically described in these regions (these have been significant resources in collecting synonyms and their metadata for Hesperomys and the MDD). Higher-level mammal taxonomy also began to take shape at this time. The publication of G.G. Simpson’s *The Principles of Classification and a Classification of Mammals* (1945), which is the foundation of modern mammalian ordinal and familial arrangements, is based on a more complete consideration of the deep time evolutionary history of both living and extinct mammals.

Collectively, these compendia helped organize mammal taxonomy following the prior peak in descriptive taxonomy, contributing to the creation of global compendia like Walker et al. (1964; and 6 subsequent editions), Morris (1965), and Anderson and Jones (1967, 1984), which provided some of the first estimates for total extant mammal species richness. Subsequently, the three editions of *A World List of Mammalian Species* (C&H1, C&H2, C&H3; Corbet and Hill 1980, 1986, 1991) and three editions of *Mammal Species of the World* (MSW1, MSW2, MSW3; Honacki et al. 1982; Wilson and Reeder 1993, 2005) provided the first complete taxonomic compendia of the modern era, each explicitly listing all contemporarily recognized extant and recently extinct mammal species.

A testament to the rapid transition from descriptive to revisionary taxonomic research is illustrated by comparing estimated mammal species totals from across this period (Fig. 1, upper panel). The earliest known estimate following the descriptive peak is ∼15,000 extant mammal species in the first edition of Storer’s *General Zoology* (Storer 1943). Although Storer does not mention how this count was determined, it can be treated as a rough estimate of the number of mammal names described as species and subspecies up to that point. We base this on the fact that Storer (1943) gave 25,000 as a combined estimate for bird species and subspecies but then reduced these numbers to 8,600 bird species and 4,400 mammal species in Storer’s (1951) second edition. Mammal species estimates further decreased to 4,237 species in Morris (1965), who based his total on various compendia plus recent primary literature that had synonymized names described in previous decades. Morris also expressed a resounding sentiment of mid-1960s mammal taxonomists:

*As more and more specimens became available and as the concept of reproductively isolated species became accepted, so the tide was turned and the “splitters”, as they were called, were replaced by the modern “lumpers”. They set about the task of cleaning up the enormous mess and sank name after meaningless name into oblivion.* (Morris 1965:18).

Estimates of total mammal species richness continued to decrease before reaching a low in 1980, when Corbet and Hill listed 4,005 recent mammal species (C&H1), being the first known publication since the peak of mammalian descriptive taxonomy to include a complete list of every then-recognized valid mammal species. C&H1 marks the shift from predominantly lumping species back to predominantly splitting species, as total estimates of extant mammal species richness began increasing thereafter (Table 1, Fig. 1).

The 20th century also saw transformative advancements in the theory and methods used to delimit species, which directly influenced mammalian systematics. Population genetics theory established the first quantitative framework for understanding the genetic variation within and between populations, providing a genetic basis for re-interpreting previously defined morphological variation (Charlesworth and Charlesworth 2017). The shift from phenetics to cladistics introduced a phylogenetic approach to analyzing shared derived traits, offering a means of reconstructing taxonomic relationships based on evolutionary theory (Hennig 1966). Understanding of the evolutionary significance of genetic heritability helped inspire formal species concepts, such as the Biological Species Concept (BSC; Dobzhansky 1937; Mayr 1942) that emphasized reproductive isolation as the basis for species limits, and the Phylogenetic Species Concept (PSC; Cracraft 1983) that diagnosed the monophyly of lineages to define species. The development of the General Lineage Concept (GLC; de Queiroz 1998) near the end of the period advocated for the definition of species as independently evolving lineages along a continuum, which helped clarify operational issues with the BSC, PSC, and other species concepts. The application of species concepts was aided by new empirical methods, including karyology, protein electrophoresis, and eventually DNA sequencing.

Molecular genetic sequencing and computational phylogenetic analysis revolutionized taxonomy in the late 20th century, resolving many longstanding questions in mammalian systematics and opening the doors to comparative genomics (O’Brien et al. 1999). The shift toward a phylogenetic and cladistic framework for taxonomy sparked the development of McKenna and Bell’s (1997) *Classification of Mammals Above the Species Level*, the first phylogeny-informed higher-level mammal taxonomy and the last publication to consider all clades of living and fossil mammals. Additionally, this period saw more taxonomically focused specimen collections, advancing knowledge of species distributions to aid in further revisionary studies (Dunnum et al. 2018). Together, these innovations enabled the modern era of mammal systematics, in which the names and taxa described in previous decades would again be revised to create more uniformly meaningful units.

### Mammalian Systematics in the Modern Era

The 44 years since the publication of C&H1 (Corbet and Hill 1980) have seen a nearly 70% increase in the number of recognized mammal species (Table 1). Similarly, total recognized species richness has increased by ∼46% in the four decades since MSW2 (Wilson and Reeder 1993; Cole et al. 1994), ∼25% since MSW3 (Wilson and Reeder 2005; Reeder et al. 2007), and ∼4% since the release of MDD1 (Burgin et al. 2018). The recent and rapid net increase in recognized mammal species is comparable to activity in other major clades of life (e.g., reptiles: +49% from 1995–2024, Uetz 2016, Uetz et al. 2024; amphibians: +60% from 1985–2024, AmphibiaWeb 2024; foraminifera: +21% from 1987–2020, Hayward et al. 2020). Parallel taxonomic efforts have focused on cataloging hyperdiverse clades (e.g., vascular plants, ∼1.3 million names for ∼0.35 million accepted species; Borsch et al. 2020) or parsing among competing taxonomic perspectives (birds: four main lists differing by ∼20% or more; Conix et al. 2024). Thus, in this section we consider recent advances in mammal taxonomy relative to two more general advances in taxonomic research: (i) new kinds of data and increased data resolution; and (ii) more sophisticated methods of species delimitation.

The shift to integrating molecular and morphological data to diagnose species limits in the late 20th and early 21st century spurred the resurgence of taxonomic study across many clades, a period regarded as the “taxonomic renaissance” (Mallet and Willmott 2003; Sites and Marshall 2003; Miller 2007). Mammals typify activities of this renaissance in two ways: (i) increased splitting of existing species through taxonomic revisions (774 species split since MSW3); and (ii) proportionately more names described as new species rather than subspecies (83.4% of available names since 2000 are described as new species rather than subspecies, versus 36.1% from 1950–1999). Despite controversy over whether this shift constitutes “taxonomic inflation” (e.g., Zachos et al. 2013; Groves 2014), the recent changes to mammal taxonomy have been bidirectional, often involving both synonymization and new species recognition. For example, 226 species-rank names that were recognized as species in MSW3 have since been synonymized, or about 1 lump for every 6 newly recognized species from 2004-2024 (considering both de novo and split names). Similarly, there has been a net increase of 123 recognized genera since MSW3, typically to establish generic monophyly, which has also been bidirectional (+178 and -55 genera). There are 26 genera with all included species described after MSW3, indicating that they were added based on the discovery of new species descriptions rather than splits. These generic revisions have changed the binomial names of at least 514 species that are recognized in both MDD2 and MSW3, a conservative estimate that does not consider generic changes to the many post-MSW3 species recognitions. Even the recognized species richness of groups such as *Microcebus* mouse lemurs, which rose from 2 to 25 species over four decades (Franz et al. 2020), was recently reduced to 19 species after an integrative taxonomic revision (van Elst et al. 2024—published after the MDD2 cutoff date).

Thus, like the preceding time period discussed above, revisionary taxonomy is perhaps the most appropriate characterization of mammal taxonomic change in this modern era (albeit with more frequent splitting than lumping).

Additional shifts from single-gene to genome-scale molecular analyses, and from linear and geometric morphometrics to computed tomographic (CT) scans, have revolutionized how species-rank units are being delimited in mammals and other taxa. The increasing use of integrative taxonomic approaches—defined as delimiting taxa via independent lines of evidence (e.g., comparing phylogeographic, morphological, ecological, and behavioral data; Dayrat 2005; Padial et al. 2010)—coupled with more rigorous statistical methods for delimiting species (reviewed in Smith and Carstens 2022) has led to revised species richness estimates within many taxa (Jörger and Schrödl 2013; Fišer et al. 2018). Documenting previously cryptic evolutionary structure has increased the informativeness of the species unit for downstream questions (e.g., biogeography, conservation; Vences et al. 2024). However, some researchers have pushed back against this movement, arguing that the recognition of “less distinct” species with histories of hybridization is a type of “taxonomic anarchy” that hampers conservation (Garnett and Christidis 2017). In response, others have emphasized that science-based taxonomy that “follows the evidence” is essential for the integrity of conservation efforts (Raposo et al. 2017; Thomson et al. 2018). Additionally, the loss of intermediate populations through human-mediated habitat fragmentation may lead to the recognition of previously connected populations as distinct species, which some argue as contributing to artificial taxonomic inflation (e.g., Clavero et al. 2024), while others express that this reflects a natural evolutionary shift that should be considered in taxonomic inferences (“speciation-by-extinction”; Seeholzer and Brumfield 2023). Establishing more formal governance structures for taxonomic decision making, at least partially mirroring the formal regulation of zoological nomenclature, is viewed by some as the path forward (Thiele et al. 2021). Others highlight the need for more pluralistic approaches, such as by developing tools to allow biodiversity databases to accommodate multiple taxonomic identifications for the same specimens and related data (Franz and Sterner 2018; Sen et al. 2021; Montoya 2022). We later discuss how this interplay among biodiversity informatic disciplines impacts decision making (see *Mammal Taxonomy in Conservation and Biodiversity Infrastructure* section below).

New data and tools have not quieted the debate regarding “how distinct” a given population should be to be recognized as a separate species, especially given the substantial histories of introgressive hybridization that are now being uncovered within some mammal clades. Throughout the 21st century, phylogenetic reconstructions based on one or a few genes have been used to justify species limits among mammals. However, it may matter more *which* rather than *how many* characters or genomic regions are shared versus distinct among putative species. That is because some mammal species can retain phenotypic and ecological distinctiveness from related species in the face of moderate to extensive histories of introgressive hybridization (e.g., mustelids, felids, bats; Li et al. 2019; Colella et al. 2021; Foley et al. 2024). Comparative genomic evidence in cats (Bredemeyer et al. 2021) and house mice (Forejt et al. 2021) suggests that some regions on the X chromosome are “protected” from recombination, causing hybrid male offspring to be sterile if altered, thereby reinforcing the distinction among lineages. In contrast, 17–93% of the genomes of European *Myotis* bats apparently have introgressive origins (Foley et al. 2024), suggesting that many genomic regions can be freely exchanged without compromising the distinctiveness of a given species.

Nevertheless, broadening such conclusions into generalities about mammals is likely premature. Many taxa lack sufficient geographic and taxonomic sampling to identify and interpret such hybridization histories (e.g., in the case of *Myotis keenii* and *M. evotis*; (Lausen et al. 2019, 2021; Morales et al. 2021; Upham et al. 2022b). Sampling difficulties are often exacerbated by the costs of fieldwork, specimen curation, and genome sequencing, but newfound efficacy sequencing genomes from preserved museum specimens promises to soon fill many populational gaps (e.g., Card et al. 2021; Raxworthy and Smith 2021; Benham and Bowie 2023). In a possible future where population genomic data are available for most mammal species, following the evidence about what genomically constitutes a different species may continue to surprise us.

Overall, trends in modern taxonomic research appear to reflect a paradigm shift in how systematists define the species unit (reviewed in Padial and De la Riva 2021). Suggested reimaginations of the subspecies rank to avoid species inflation (Hillis 2020; Dufresnes et al. 2023) are balanced against the uneven history of subspecific classification that may render these categories ambiguous (Patton and Conroy 2017; Burbrink et al. 2022). For now, the MDD has refrained from formally recognizing subspecies in the taxonomy of mammals but may start doing so in some groups (e.g., bats) where species-to-subspecies relationships have been extensively tracked and thus retain meaning (Simmons and Cirranello 2024). More broadly, it is remarkable that the once-vigorous philosophical debate surrounding the validity of different species concepts is now largely superseded by integrative taxonomic approaches. We acknowledge that the historical discourse is important for understanding the current state of mammal systematics, but we defer to other sources that discuss it in depth (see Bradley and Baker 2001; Gippoliti and Groves 2012; Gippoliti et al. 2013, 2018; Zachos and Lovari 2013; Zachos et al. 2013, 2020; Zachos 2015, 2018a, 2018b; Gippoliti 2019; Taylor et al. 2019). Contemporary mammal systematics now focuses on evaluating how many independent lines of evidence simultaneously support a taxon’s distinctiveness (i.e., integrative taxonomy; Dayrat 2005; Padial and De la Riva 2021). To this end, recent efforts have sought to establish a “traffic-light system” to evaluate levels of taxonomic support for felid species (Kitchener et al. 2022), which parallels a proposed “scoring system” for bird species delimitation (Tobias et al. 2010, 2021). This shift toward integrative taxonomy as opposed to choosing a single species concept underscores the importance of sharing key information in up-to-date taxonomic databases.

Considering these trends, the relevance of integrating the latest evidence regarding mammal species boundaries into biological research and conservation efforts worldwide is only expected to grow.

### Patterns and Shortfalls in Global Mammal Biodiversity

Despite recent progress, considerable gaps remain in our collective knowledge of mammal species-rank taxonomy and geographic distributions—termed the Linnaean and Wallacean shortfalls, respectively (Hortal et al. 2015). These shortfalls pose hurdles to zoonotic disease and other biomedical research (Upham et al. 2021) as well as to the implementation of species conservation and management plans (Senior et al. 2024). This is particularly true for mammals given that they often serve as sentinels of change in threatened ecosystems (Moore 2008; Marneweck et al. 2022) or key nodes in zoonotic disease networks (Plowright et al. 2021; Venkatesan 2023; Weber et al. 2023). Through the compilation and curation of up-to-date taxonomic, geographic, and nomenclatural data, the MDD aims to continue lessening these shortfalls for mammalogical research. This curatorial work is synergistic with two parallel efforts in the biodiversity research community: (i) at the specimen level, where holistic specimens (Cook et al. 2016) and their derived data are becoming linked to form “digitally extended specimens” (Lendemer et al. 2020; NAS 2020); and (ii) at the publication level, where related data from publications are being extracted and linked to form “digitally accessible knowledge” (Fawcett et al. 2022). The non-copyrightability of primary research findings, particularly of the parts of articles that describe how names are conceptually applied to taxa (i.e., taxonomic treatments; Agosti and Egloff 2009; Benichou et al. 2023), provides an explicit legal framework for continuing to integrate these kinds of liberated and linked biodiversity data into the MDD.

Efforts to study the Linnaean shortfall typically estimate how many undescribed species likely exist and how long until the gap will be filled, using estimates for current rates of description based on authority years of species epithets. Using this approach in 2018, the MDD team estimated that recognized extant and recently extinct mammal richness will increase to 7,409 species by 2050 and to 9,009 by 2100 if observed trends in historical species descriptions were to continue (Burgin et al. 2018). That pace exceeded the previous estimate of 7,500 total species based on the same method (Reeder et al. 2007). However, those estimates were both based on partial histories of mammal species description that ignore taxonomic changes resulting from species splitting and lumping, instead only focusing on de novo species descriptions. Lessa et al. (2024) reviewed this problem in detail, concluding that “relevant estimates of the number of known and unknown species will only be achieved by accounting for the dynamic nature of the taxonomic process itself” (p. 1365). Indeed, if we reconsider the predictions of Burgin et al. (2018) using the same approach, the expectation for 418 valid de novo species descriptions from 2010-2019 compares to only 369 currently valid species that were actually described de novo (i.e., with an authority year during this interval). That decadal total compares to 315 during 2000-2009 (originally 341 species in Burgin et al. 2018 but reduced by synonymizations in MDD2). Thus, these data reinforce the perspective that rates of de novo species description are only part of the “discovery rates” story, with the other major component being rates of species splitting and lumping that result from revisionary taxonomic efforts.

To more fully investigate this dynamic for mammals, we calculated the rate of new species *recognition* based on the estimated species totals reported in authoritative taxonomic compendia through time. These totals each represent the net outcome of species splits, lumps, and de novo descriptions at a given time point. Conducting a linear regression of mammal species totals sourced from 14 taxonomic compendia over 44 years (1980 to 2024; Table 1, Fig. 1 upper panel) reveals a rate of 64.9 species recognized per year, which is higher than the rate of 27.4 species/year reported in Burgin et al. (2018) based on de novo descriptions alone. If this trend continues, we estimate that the total recognized mammal species richness will increase to ∼7,084 species by 2030 and ∼8,382 species by 2050. Both of these estimates are considerably higher than those given by Burgin et al. (2018) and Reeder et al. (2007) but are nonetheless realistic owing to the combined consideration of species splits, lumps, and de novo descriptions. The estimate of ∼11,625 species by 2100 seems more far-fetched given that this trend is likely to level off at some point, but the ∼70% increase over the current MDD2 total would be equivalent to the 1980–2024 increase from C&H1.

The potential for unidentified cryptic mammal species remains high, particularly within small-bodied and wide-ranging taxa. Parsons et al. (2022) used mitochondrial genetic data from a subset of 4,310 mammal species to find which covariates best predict unrecognized cryptic species, with body mass and range size ranking the highest. A meta-analysis across metazoans similarly supported this conclusion (Cahill et al. 2024). Unsurprisingly, rodents, bats, and eulipotyphlans are predicted to be the largest reservoirs of unrecognized cryptic mammal species despite the high rate of new species recognition in these clades. However, less specious mammal taxa like Hyracoidea, Scandentia, Pilosa, Pholidota, Lagomorpha, and Perissodactyla that have received comparatively less taxonomic attention in the 21st century are also more likely to contain cryptic species (Parsons et al. 2022). Within Pholidota, for example, a potentially new species was identified from DNA derived from tissue confiscated from the illegal pangolin trade (Gu et al. 2023), and another new species was also recently described (described as *Manis indoburmanica*, but *Manis aurita* Hodgson, 1836 may have nomenclatural priority; Wangmo et al. 2025). Similarly, a new species of Hyracoidea was described with evidence that further cryptic hyrax species exist (*Dendrohyrax interfluvialis*; Oates et al. 2022). Didelphimorphia stands out as having both a high degree of potential for cryptic species and extensive recent taxonomic research, with over 50 new species recognized since MSW3 (39.7% of valid Didelphimorphia species in MDD2). Conversely, the order Primates likely has considerably less remaining cryptic species, owing to focused and ongoing taxonomic research during the 21st century (161 new species since MSW3, 30.8% of valid Primates species in MDD2). Clade-specific taxonomic research thus remains imperative across the mammalian phylogeny, even in groups like Perissodactyla and Artiodactyla that are historically thought to be well-studied but still mostly lack integrative taxonomic revisions.

Geographically, recent progress in bridging the Linnaean shortfall for mammals has followed a latitudinal gradient, with more newly split species and de novo descriptions coming from tropical regions (Fig. 2, Table 3). The Neotropics and Indomalaya have particularly high proportions of newly recognized mammal species since MSW3, including 9 of the top 10 most new species-rich countries (Fig. 2A and 4B) and similarly at the continental scale (South America and Asia; Fig. 2B). Only around half of these newly recognized tropical species are splits, as opposed to in the Nearctic/North America and the Palearctic/Europe where split species form a higher proportion of new species recognitions since MSW3. These geographic patterns of taxonomic activity are consistent with previous estimates (Patterson 2000; Reeder et al. 2007; Burgin et al. 2018; Moura and Jetz 2021). In contrast, some geographic regions have fewer newly recognized species than expected given their latitudinal location, habitat type, and total species richness. This is most evident in parts of Africa and Asia, such as West and Central Africa, Central Asia, most of India, and some Indonesian islands, supporting calls for taxonomic work in those regions (e.g., Klopper et al. 2002; Chatterjee et al. 2020; Monadjem et al. 2024). In terms of new species descriptions over the 21st century (Figs. 3), de novo species descriptions appear to be most concentrated in areas of high species endemism, such as mountains (e.g., Andes, Himalayas, East African Highlands), islands (e.g., Madagascar, Philippines, Indonesia), and other biodiversity hotspots (e.g., Atlantic Forest). However, some of these patterns likely represent regional biases in taxonomic research and museum specimen collection. India, the Indonesian islands of Borneo, Sumatra, and Java, and West Africa have comparatively low rates of de novo species description but include large land areas with diverse habitat types conducive to endemism. The regions discussed above also appear to represent “biodiversity blindspots” according to a recent global assessment of digitized records in natural history collections (Ball et al. 2025), further highlighting the need for both specimen collection and digitization in these regions.

The Wallacean shortfall of incomplete geographic knowledge pertains to both newly described and previously recognized mammal species. As with earlier eras of mammal species descriptions (e.g., the peak of descriptive taxonomy in the early 19th century), some of the newly described species, particularly those known from fewer specimens, are likely to be synonymized in the future as additional populations and characters are sampled (Alroy 2002). Natural history collections and biorepositories play a central role in bridging geographic shortfalls by maintaining the core collections of voucher specimens used to document species distributions over time and space (Bakker et al. 2020). However, historical biases in mammal surveys have left large sampling gaps in tropical regions despite their high biodiversity and rates of recent species description (Fig. 4A, 4B). In addition, most specimens from those regions are housed in Europe and North America (see figure 2 in Heberling et al. 2021). This imbalance between where mammals live and where their specimens are preserved helps to perpetuate geographic disparities in biodiversity knowledge and conservation outcomes (see Pritchard et al. 2022). Thus, investing in resources like the MDD that digitally link and distribute this knowledge is a key step toward expanding access to research materials in highly biodiverse regions.

### Mammal Taxonomy in Conservation and Biodiversity Infrastructure

Despite advances in the understanding of species-rank taxonomy and geographic distributions since MSW3, ∼25% of wild mammal species are ‘understudied’ by the IUCN Red List — either Not Evaluated (i.e., unclassified; ∼14%) or classified as Data Deficient (∼11%; Fig. 6). The IUCN Red List has assessed the conservation status of 5,983 mammal species as of version 2024-1, which is only 494 species more than were assessed in the 2008 IUCN Red List version (5,489 species; summarized in Schipper et al. 2008). The total number of IUCN-assessed species is also 759 fewer than recognized in MDD2 (excluding domestic species not assessed by the IUCN). This IUCN-to-MDD2 gap is due to multiple factors, both infrastructural and sociological, that lead to delays in the integration of the latest research findings into the IUCN taxonomy.

Unfortunately, delayed IUCN assessments are not unique to mammals and are viewed as a top priority for improving conservation outcomes globally (reviewed in Bachman et al. 2019). This lag in taxonomic change integration is also not unique to the IUCN, reflecting other ongoing challenges in biodiversity data science related to the linking and dissemination of primary research findings (e.g., Lendemer et al. 2020; Upham et al. 2021; Benichou et al. 2023; Sterner et al. 2023). The MDD is working to make progress in this domain via the curation and regular versioning of the collective implications of recent taxonomic publications. However, there is potential for much closer integration of MDD efforts with downstream biodiversity resources like the IUCN Red List. Failing to do so risks unnoticed species declines or population extinctions, particularly for the ∼25% of mammal species that are understudied. Untracked taxonomic rearrangements also risk the misallocation of resources (e.g., for species like *Sylvilagus mansuetus* that the IUCN lists as CR but is not recognized in MDD2 due to synonymization with *S. bachmani*; Álvarez-Castañeda and Lorenzo 2016). Thus, greater MDD-to-IUCN integration carries potential for both speeding up processes of assessment and improving outcomes in mammal conservation.

Conservation practices in mammals often prioritize named taxonomic units like species and subspecies as the primary units of conservation, as seen in global frameworks like the IUCN Red List and national frameworks like the United States Endangered Species Act and Canadian Species at Risk Act. However, this practice can overlook finer-scale intraspecific diversity (Des Roches et al. 2021) and taxonomically unrecognized cryptic species in understudied taxa (i.e., “dark extinctions”; Boehm and Cronk 2021), resulting in the loss of cryptic biodiversity (see review on cryptic species conservation by Hending 2024). To address these issues, some researchers and managers have argued for formally recognizing intraspecific variation as subspecies (e.g., Dufresnes et al. 2023) or non-Linnaean conservation units (e.g., Evolutionarily Significant Units, ESUs; Casacci et al. 2014). However, others have pointed out that although some conservation organizations recognize subspecies and other infraspecific conservation units (e.g., the IUCN and United States Endangered Species Act), the species rank has remained the primary “currency of conservation” in various mammal clades (Rylands and Mittermeier 2014; Gippoliti et al. 2018). Ideally, all units along the population-to-species continuum would receive some conservation protections regardless of their taxonomic rank, but the finite resources available for conservation and the incomplete assessments of global biodiversity have prevented this reality (Gippoliti and Amori 2007; Coates et al. 2018). As a result, some researchers have used conservation priorities to guide taxonomic practices rather than vice versa, leading either to the splitting of species into multiple smaller conservation units or the lumping of species to conserve those same units as subspecies or ESUs. The relationship between taxonomy and conservation is dynamic, and debate over the extent to which conservation concerns should influence the science of taxonomy will last long into the future.

The recent taxonomic history of primates and terrestrial ungulates (Perissodactyla and non-cetacean Artiodactyla) best illustrates the impact that conservation can have on species recognition. Both groups have been at the center of splitting versus lumping debates (e.g., Rylands and Mittermeier 2014; Gippoliti et al. 2018) and both groups include many traditionally charismatic mammal species that are threatened with extinction (∼64% of primates and ∼40% of wild ungulates in MDD2 are listed as threatened by the IUCN). Both groups also have had their taxonomies updated more regularly by the IUCN than other groups of mammals (only ∼3% of primates and ∼8% of wild ungulates in MDD2 are Not Evaluated by the IUCN). However, the similarities end there, as these two groups have experienced considerably different taxonomic trajectories thus far in the 21st century. Primates experienced more species splitting starting with the publication of Groves’ (2001) *Primate Taxonomy*, which was controversial at the time but initiated a wave of taxonomic revisions that massively increased the recognized primate species richness (176 species in Napier and Napier 1967 versus 376 in MSW3 and growing to 522 in MDD2; see Groves 2014). The publication of Groves and Grubb’s (2011) *Ungulate Taxonomy*, which split many ungulate species, served as a similar catalyst for taxonomic revision within ungulates. However, the outcome of that major shakeup in ungulate taxonomy did not have the same effect as in primates, since later authorities (including the MDD after version 1.2) have generally reverted to pre-2011 taxonomies and focused on making taxonomic changes based on more integrative taxonomic revisions (see *Taxonomic Decision-Making* section of Methods for further details). For example, recognized *Giraffa* diversity has ranged from 1–11 species, but most workers now recognize 4 species based on integrative taxonomic approaches (Winter et al. 2018; Petzold and Hassanin 2020; Coimbra et al. 2021; Kargopoulos et al. 2024). Both primates and ungulates require further taxonomic research, but the non-uniformity of taxonomic ideologies being applied remains a hotly debated topic (e.g., Taylor et al. 2019; Zachos et al. 2020).

The MDD taxonomy increasingly serves as an arbiter of species recognition for conservation organizations at global (e.g., IUCN), national (e.g., NatureServe), and regional levels (e.g., state agencies). Additionally, the MDD aims to serve as a foundational taxonomic connection between other biodiversity resources, such as species record aggregators (e.g., GBIF, Map of Life, iNaturalist), digitized museum archives (e.g., Arctos, Symbiota, iDigBio), and various other biological databases (e.g., GenBank, NCBI Virus, FuTRES, Morphosource, Global Biotic Interactions—GloBI). Efficient communication and application of biodiversity knowledge between conservation and biodiversity resources requires agreed-upon organismal units that can serve as foundational designations for conservation decisions made on those units. Most often these units are species (but specimens or individuals are also commonly used). The value of accurately communicating ‘species’ categories highlights the basal importance of taxonomy and nomenclature in biodiversity infrastructure (Grieneisen et al. 2014; Sterner et al. 2020a).

However, as discussed throughout this article, taxonomy and nomenclature are by no means static—the creation of species lists is plagued by gaps in biodiversity knowledge (i.e., the Linnean and Wallacean shortfalls), differing taxonomic opinions, failure to track and digitally integrate those changing taxonomies, and inconsistencies in how the ICZN Code is implemented (Garnett and Christidis 2017; Raposo et al. 2017, 2021; Franz and Sterner 2018; Upham et al. 2021). Nevertheless, the value of updated species lists to biodiversity science outweighs the pitfalls of maintaining them. The value of species-centric taxonomic databases like the MDD grows in proportion to the number of disparate sources of biodiversity knowledge they can effectively integrate. Doing so improves the rigor of a wide range of studies in conservation, biodiversity, and biomedical sciences that rely on accurate taxonomy, facilitating work that might otherwise go undone.

### Governance

Moving forward, the MDD is in the process of forming external taxonomic subcommittees that are scheduled to begin activity in early 2026. Subcommittee members will include global representation of taxon-specific experts to help with the ongoing process of curating mammal taxonomic, geographic, and nomenclatural information, serving an advisory role to the ASM’s Biodiversity Committee in coordinating the MDD. Initial plans have divided mammal clades across 25 subcommittees, each one focusing on a clade or composite taxonomic group ranging in size from 18 species (Perissodactyla) to over 800 species (Cricetidae, Muridae). Each subcommittee will have members shared with one or more IUCN Species Survival Commission Specialist Group, of which there are currently 36 for mammals (IUCN 2024b), aiming to facilitate direct coordination between MDD taxonomic changes and the IUCN Red List assessment process. MDD taxonomic subcommittees will also be encouraged to formally issue ‘subjective decisions’ in cases of taxonomic conflict in the primary literature, following the model of Zenodo self-publishing previously established for bats (see *Taxonomic Decision-Making* section in the Methods—Francis et al. 2022, 2023; Upham et al. 2022b). A key goal of subcommittee efforts is to organize archives that contain full citations and textual links to all the publications used to justify taxonomic decisions in the MDD. These archives can then be directly shared with outside research groups to lower barriers to access and catalyze further research. A pilot of this approach is available on https://batlit.org, containing >20,000 publications pertaining to bat taxonomy and ecology that are a mix of open and closed access works uploaded to Zenodo for controlled sharing with collaborators (Sherman et al. 2024). Expanding this pilot effort to encompass publications relevant to taxonomic decisions across Mammalia would parallel similar initiatives in drosophilid flies (Bächli 2024) and reptiles (Uetz et al. 2023). Collectively, the formation of MDD taxonomic subcommittees aims to create a structured community of experts for sharing literature and curating taxonomic knowledge of mammals globally. ASM membership will not be required for inclusion, which underscores the expected global breadth of participant expertise as well as the need to effectively coordinate subcommittee activities using online collaboration tools.

Overall, the MDD team currently strives to “follow the science” and update the MDD based on peer-reviewed scientific articles as they are published and with as few subjective decisions as possible. However, this can result in taxonomic revisions that are ricochetal, bouncing between article-and-response cycles (e.g., recent back-and-forth debates concerning the bat genus *Lasiurus*; Baird et al. 2015, 2017, 2021; Ziegler et al. 2016; Novaes et al. 2018; Teta 2019). It may also result in revisions that the broader mammalogical community see as weakly corroborated. With these limitations in mind, we recommend that conservation authorities and other biodiversity databases use the MDD as a starting point to build species lists for policy creation and as a backbone for inter-database connectivity. Soon, the MDD team aims to work with the taxonomic subcommittees to establish a multi-dimensional ‘grading system’ that will summarize information on both the *extent* and *quality* of corroborating taxonomic evidence for particular species. This grading system will expand on Kitchener et al.’s (2022) proposed ‘traffic-light’ categories by establishing a continuous rubric for grading the completeness of data sampling and statistical strength of that taxonomic evidence. The number of independent datasets that support species recognition (e.g., genetic, ecological, morphological) would still be considered, but the new quality rubric will add degrees of nuance that are hidden within the traffic-light system’s categories (green, yellow, red, and black—only one of which is “go”).

Building this grading system will provide a more flexible means of synthesizing the extent to which integrative taxonomic evidence supports a given species as being distinct from others. Once established, the contained evidence of this grading system will allow researchers, conservationists, and policymakers to compare support and identify gaps in knowledge of different taxa. This will, in turn, enable better-informed decision making, including in cases where an organization chooses to deviate from the recognized MDD taxonomy. Enabling a plurality of taxonomic viewpoints rather than enforcing a single one is a feature that we contend will maximize the equitability of biodiversity outcomes (see reviews by Franz and Sterner 2018; Montoya 2022; Sterner et al. 2023).

### Future Directions and Conclusion

Continued improvements to the MDD are targeted in three areas: (i) curating new and existing data sources; (ii) linking those curated data to other public databases; and (iii) informing future curation using those newly linked resources (Fig. 7). This iterative process will help establish the MDD as an open-access portal for mammalogical research globally. We aim for each MDD species page to eventually serve as the primary hub for connecting researchers to external biodiversity information about that mammal species. Much of this content will be hosted on other websites indexed using versions of the MDD taxonomy. Databases such as the Integrated Taxonomic Information System (ITIS 2025) and Catalog of Life (CoL 2025) are increasingly critical for national and international processes of biodiversity decision-making, but they still primarily rely on MSW3-based taxonomies and synonymies for mammals. The MDD’s continuously curated data now presents an excellent opportunity for greater cross-linking among various databases that contain mammal biodiversity information. National Center for Biotechnology Information (NCBI) resources like GenBank, RefSeq, and NCBI Virus will especially benefit from greater cross-linking of deposited genetic data with updated taxonomies for mammals and other taxa (Garg et al. 2019). Currently, only record authors or administrative batch edits can update NCBI taxonomy (Schoch et al. 2020), but if the MDD team can identify cases where taxonomic updates are unambiguous, then feeding batch recommendations to NCBI curators could allow for greater integration.

**Figure 7.**
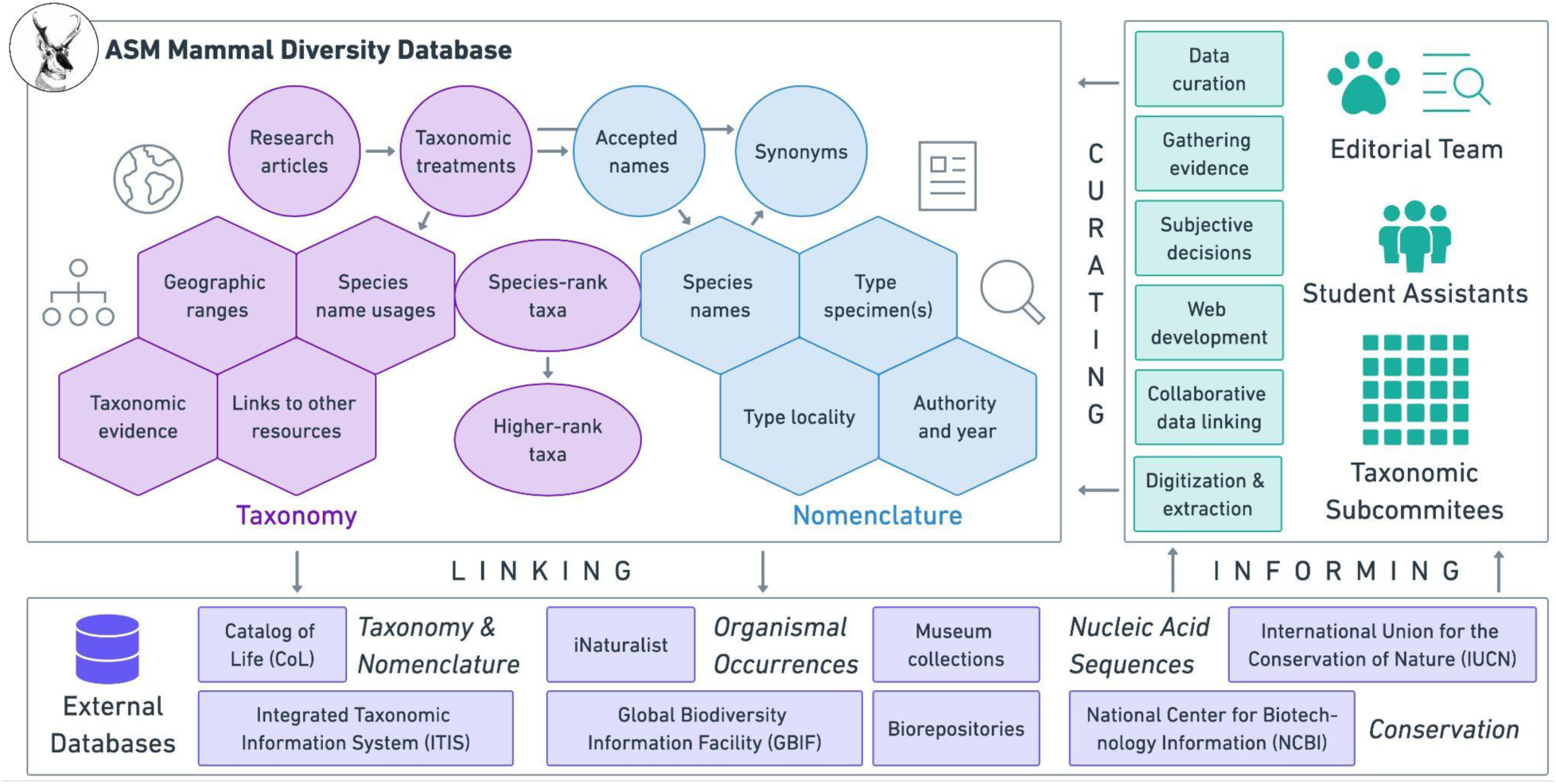
Overview of the MDD’s structure and governance for future versions. Taxonomic knowledge will continue being extracted from research articles, each of which presents peer-reviewed evidence in support of a perspective on the limits of species- and higher-rank taxa. Nomenclatural information in those taxonomic treatments will also be curated, including the status of available names for a given taxon and the associated physical evidence for those names in the form of one or more type specimens. The MDD editorial team, consisting of both volunteers and paid student assistants, will be advised by new sets of taxonomic subcommittees that will help gather and curate data while providing expert guidance in cases where subjective decisions need to be issued. Linking of the resulting MDD data with external databases will remain a top priority, both to provide up-to-date information to those sources and to inform the taxonomic curation process itself.

Considering MDD plans relative to broader activities of the American Society of Mammalogists (ASM), which generously supports the MDD through the Biodiversity Committee, there is potential for greater taxonomy-related coordination across Society efforts. Leaders of the Public Education Committee that curates a US state list of mammals and the Mammal Images Library (MIL) Committee that curates expert-identified photographs of mammals both regularly communicate with the MDD team, but the activities of those committees are not directly integrated with MDD taxonomic updates. For example, hosting MIL images on MDD species pages may help flag needed naming updates, enabling the maintenance of the MIL’s current hand-harmonization with the MDD taxonomy. Similarly, the long-time ASM publication *Mammalian Species* (*MS*; 1969-present) may benefit from being directly indexed using the MDD taxonomy, which would help flag *MS* species accounts needing updates after taxonomic changes. Lastly, MDD efforts could help address ethical concerns over the use of eponyms in mammalogy (i.e., species named after people). Recent discussions over whether eponyms should be changed regardless of whether they were named after controversial figures do not have easy solutions (e.g., DuBay et al. 2020; Guedes et al. 2023; Thiele 2023; Winker 2024).

Wholesale changes to scientific names are untenable without more sophisticated biodiversity infrastructure for renaming millions of species observations (Ceríaco et al. 2023). However, in mammals at least, eponymous *common names* would be much easier to begin changing given that they are unregulated by any governing body (in contrast to birds; Winker 2024). As a result, Chen-Kraus et al. (2021) recently proposed changing the eponymous common names of 140 primate species. Clearly there is a need to seriously consider such proposals, and the MDD could function as a community-based platform to help reach consensus on this and other controversial issues.

In conclusion, MDD2 and future versions offer a platform for the community curation of often hard-to-access but fundamentally important data that underpins biodiversity knowledge of living and recently extinct mammals. With 6,759 recognized mammal species and >50,000 species-rank names, MDD2 provides a comprehensive resource that not only lessens aspects of the Linnean and Wallacean shortfalls but also highlights critical remaining knowledge gaps. The hard-won nomenclatural metadata included in MDD2 provide researchers with the foundational tools needed to conduct taxonomic revisions, resolve nomenclatural conflicts, and refine species boundaries. By ensuring that these metadata are linked to broader biodiversity infrastructure, the MDD supports more robust and integrative approaches to mammalian taxonomy and conservation planning than would otherwise be possible. MDD2 also reveals a pressing conservation challenge: the ∼25% of mammal species that remain understudied on the IUCN Red List raises alarms about the need to address these knowledge gaps and prevent unrecognized biodiversity loss (Ceballos et al. 2017). By establishing taxonomic subcommittees and fostering global collaborations, the MDD is positioned to continue providing authoritative taxonomic updates that advance both scientific and conservation priorities. The MDD and parallel databasing efforts across the Tree of Life underscore the general recognition across the biological sciences that high-quality curated information regarding how species are classified (taxonomy), how names are applied (nomenclature), and where those species live (biogeography) is critical for the accuracy of downstream research. Species, their natural histories, and our scholarly history of investigating those boundaries are key for advancing humanity’s collective knowledge of biodiversity.

## ACKNOWLEDGEMENTS

We would like to thank the American Society of Mammalogists for hosting, supporting, and funding the MDD and its student workers, as well as the NSF VertLife Terrestrial grant (DEB: 1441737) for initial logistical support for this project and NIH 1R21AI164268 and 1R35GM156919 to NSU for continued development. We are grateful to the ASM Biodiversity Committee for advice, support, and input on the creation and curation of the MDD, in particular Bruce D. Patterson and DeeAnn M. Reeder for generously encouraging the establishment and persistence of this effort. We would also like to thank the IUCN Global Bat Taxonomy Working Group for input and collaboration on the taxonomy and nomenclature of bats, highlighting key contributions from the primary curators of the Bats of the World database, Nancy B. Simmons and Andrea L. Cirranello. We thank Joseph A. Cook and the Cook Molecular Evolution and Ecology Lab at the University of New Mexico for critical edits and comments on this manuscript. Lastly, we would like to thank the global mammalogical community as a whole and the natural history collections community in particular for their collective efforts to document mammal diversity, as well as continued support, feedback, and diligence in sending the MDD team publications to incorporate into the MDD.

## DATA AVAILABILITY

The R code used to analyze these data, plot figures, and create supplementary files is available in a GitHub repository accessible at https://github.com/mammaldiversity/MDDv2_ms.

## SUPPLEMENTARY DATA

Supplementary Data SD1.—MDD2 species list, plus associated metadata, in a zipped folder.

Supplementary Data SD2.—MDD2 synonym list, plus associated metadata, in a zipped folder.

Supplementary Data SD3.—All MDD version species lists, plus associated metadata, in a zipped folder.

Supplementary Data SD4.—All MDD ‘diff’ files, plus associated metadata, in a zipped folder.

Supplementary Data SD5.— ‘Diff’ file specifically between MDD1 and MDD2, plus associated metadata, in a zipped folder.

Supplementary Data SD6.—Summary of nomenclatural data and species totals by taxonomic group, plus associated metadata, in a zipped folder.

Supplementary Data SD7.—Summary of nomenclatural data and species totals by geographic region, plus associated metadata, in a zipped folder.

Supplementary Data SD8.—Summary of nomenclatural data totals by nomenclatural and validity status, plus associated metadata, in a zipped folder.

Supplementary Data SD9.—Total available and valid names described per year since 1758 and summary of 10-year running means used to create Figure 1, plus associated metadata, in a zipped folder.

Supplementary Data SD10.—Occurrence matrix for species country and offshore territory distribution data in a zipped folder.

Supplementary Data SD11.—Per country usage of the MDD website at mammaldiversity.org in a zipped folder.

## Box 1.

Definitions for taxonomic and nomenclatural terms relevant to this publication.

### Definitions Box

**Taxonomy** - the science of naming, defining, and classifying organisms into taxonomic entities, dealing with both species- (**Alpha Taxonomy**; species, subspecies) and higher-rank taxonomic groups (**Beta Taxonomy**; genus, family, order, etc.); taxonomic changes result in changes to biological concepts, by lumping taxa together (**Lump**), splitting a taxon into multiple taxa (**Split**), descriptions of new names for previously undescribed taxa (**De Novo**), or moving taxa between different higher taxa (e.g., species moving between genera).

**Taxon/Taxa** - a set of organisms together forming a single defined biological grouping at any taxonomic rank, with each ranking nested within the rank above; **Species-rank Taxa** include species, subspecies, and formerly used infrasubspecific rank names (e.g., forms, varieties); **Higher-rank Taxa** include all taxonomic rankings above the species-rank (class to subgenus applicable for the MDD).

**Nomenclature** - the process of assigning scientific names to taxa based on a set of standardized rules in which each species has a two part name, starting with a genus and ending in a specific epithet (**Binomial Nomenclature**); nomenclatural changes result in changes to the name used for a taxon that do not change its taxonomic concept; zoological nomenclature for species-, genus-, and family-rank names are governed by the rules published in the **ICZN Code** (ICZN 1999), written by the International Commission on Zoological Nomenclature (ICZN).

**Authority** - the surnames of the author(s) of the original description of a scientific name followed by the year that name was described; for species-rank names, parentheses are added around the entire authority when a specific epithet was described in a different genus than it currently resides; the year listed in the authority should be the year the name was published according to the ICZN Code (although the year of description does not always match what is printed on the publication itself).

**Name-bearing Type** - a biological entity that is permanently attached to a scientific name, acting as a reference to identify which taxon a scientific name belongs to; for species-rank names, the type is usually a single animal, some part of which or its entirety is usually preserved as a museum specimen (**Holotype**), or two or more animals preserved as museum specimens (**Syntypes**); syntypes can be later restricted to a single type specimen within the series as needed (**Lectotype**); when the original type material for a species-rank name has been lost or destroyed, one or more specimens should be designated as a **Neotype**; family-rank names have a **Type Genus** and genus-rank names have a **Type Species**, which are not based on physical biological materials.

**Type Locality** - the geographic locality from which the type material for a species-rank name originated, which may be later restricted or redefined if necessary.

**Valid Name/Taxon** - a scientific name that represents the correct name for a biological taxon as defined by the rules in the ICZN Code and recognized by a particular authority; usually but not always the oldest available name.

**Synonym** - a scientific name that has been identified as representing a specific taxon, but is not the valid name for that taxon; synonyms can be names initially meant to represent a new taxon (both Available and Unavailable Names for nomenclatural use, as defined below), or may represent later use of those names (Name Combinations or Spelling Variants; see definitions of these terms below).

**Available Name** - a scientific name that meets the standards for nomenclatural availability as defined by the ICZN Code; some names may be available for nomenclatural use but are not assignable to a single species, such as a **Nomen Dubium/Nomina Dubia** (a name completely unidentifiable to a valid taxon), **Species Inquirenda/ae** (a name not currently identified to a valid taxon, but with the potential to be), or names based on a **Composite** (type material from multiple individuals of different valid species) or **Hybrid** type material (type specimen of interspecific hybrid origin).

**Unavailable Name** - a scientific name that does not meet the standards for nomenclatural availability according to the ICZN Code, often resulting from the name being improperly published (**Unpublished Name**) or published without a taxonomic description (**Nomen Nudum/Nomina Nuda**), among other reasons.

**Preoccupied Name** - an otherwise available scientific name that matches the spelling of an older name within the same taxonomic rank, making it unavailable for nomenclatural use; in species-rank names, the preoccupying name can either be originally described in the same genus (**Primary Homonym**), or subsequently moved into that genus (**Secondary Homonym**); when the name for a currently recognized species is preoccupied, a **Nomen Novum/Nomina Nova** (new name) should be described to replace the preoccupied name.

**Name Combination** - the unique combination of a genus and specific epithet (or subspecific epithet) to form the scientific name of a then-valid species or subspecies (and forms or varieties in older literature); name combinations are synonymous names that are not currently recognized because they are not type-bearing or valid, but they can represent the history of taxonomic change associated with a name across genera, species, and subspecies.

**Spelling Variant** - a difference in spelling from a scientific name’s original spelling, which can be incorrect in the original publication (**Incorrect Original Spelling**) or later publications (**Incorrect Subsequent Spelling**), or a purposeful change considered to be unjustified (**Unjustified Emendation**) or justified (**Justified Emendation**) according to the ICZN Code; justified spelling changes are usually those of specific epithets with adjective etymological roots, which change to match the etymological gender of the genus, or to Latinized names with non-standard characters (e.g., ñ, ö), although there are other rare exceptions allowing for justified emendations to a name’s original spelling.

